# Intraspecific sequence variation and complete genomes refine the identification of rapidly evolved regions in humans

**DOI:** 10.1101/2025.10.20.683446

**Authors:** Yanting Luo, Riley J. Mangan, Seth Weaver, Sarah A. Zhao, Federica Mosti, Michael C. Thomas, Ravi Karra, Debra L. Silver, Manolis Kellis, Craig B. Lowe

## Abstract

Summary

Humans exhibit significant phenotypic differences from other great apes, yet pinpointing the underlying genetic changes has been limited by incomplete reference genomes and a reliance on single assemblies to represent a species. We aligned 20 telomere-to-telomere (T2T) assemblies spanning great ape divergence and variation to define 1,596 Consensus HAQERs (consensus human ancestor quickly evolved regions), regions that diverged rapidly between the human-chimpanzee ancestor and an ancestral node of modern humans. Unlike prior HAQER sets, Consensus HAQERs incorporate population variation, reducing the likelihood of intraspecies variation appearing as interspecies divergence. Consensus HAQERs exhibit signatures of elevated mutation rates, ancient positive selection, bivalent regulatory function, are enriched in disease-linked loci, and often emerged in previously inaccessible repetitive DNA. Through multiplex, single-cell enhancer assays, we identify HAQERs as active enhancers in the developing brain and cardiomyocytes, highlighting their potential contributions to human-specific gene regulation across multiple tissues.

**Highlights:**

● Telomere-to-telomere alignments of diverse human and great ape genomes identify 1,596 Consensus HAQERs, regions of rapid sequence divergence separating human ancestors from other great apes.
● Consensus HAQERs exhibit signatures of both elevated mutation rates and ancient positive selection.
● HAQERs are enriched in bivalent regulatory elements and disease-linked loci.
● Multiplex, single-cell gene regulatory assays identify HAQERs as enhancers in the developing heart and brain.

## Introduction

Humans have undergone dramatic phenotypic evolution since our split from a common ancestor with chimpanzees, including cortical expansion^1^, enhanced cognition^2,3^, and human-specific disease susceptibilities for neurodegeneration^4^ and neuropsychiatric disorders^5,6^. Of the tens of millions of mutations separating humans and chimpanzees^7^, it is likely that only a small fraction make significant contributions to human-specific traits^8^, making it challenging to uncover the genetic architecture that underlies phenotypic differences. Searches for functionally important mutations in human evolution have focused not only on exons^9–12^, but also on noncoding gene regulatory elements such as enhancers and promoters^13–15^ due to the central role of gene regulation in both evolution^16^ and disease^17^. This approach is exemplified by human accelerated regions (HARs), elements highly conserved across mammals but rapidly diverged in humans^18–24^. These regions represent human-specific modifications of existing functional elements, yet it remains unclear whether morphological changes across lineages primarily result from the modification of existing elements or the forging of new functional sequences^25,26^.

To better understand the potential of new functional elements, we previously defined human ancestor quickly evolved regions (HAQERs) as the most divergent regions of the human genome without preconditioning on prior negative selection. We found that many HAQERs act as *de novo* enhancers forged from previously neutral DNA^27^. This discovery is consistent with theories that humans may have a number of recently-gained neurodevelopmental gene regulatory elements^28^. Recent work has associated HAQERs with signatures of ongoing negative selection^29^, gene regulation during human cortical development,^30^ and human language^31,32^. Repetitive and complex regions of the genome, including telomeres, centromeres, and segmental duplications, are often absent from “gapped” reference assemblies. This is the case for hg38, where HAQERs were originally ascertained, which is limiting for comparative analyses of human-specific divergence. Advances in long-read sequencing and assembly algorithms have ushered in the era of complete telomere-to-telomere (T2T) genomes, resolving the ∼8% of the human genome missing from prior references^33,34^. Using these assemblies, we generated a T2T HAQER set that expanded the catalog of human-specific rapidly evolved regions, finding HAQERs to be enriched in previously unassembled sequences^32^.

Prior HAQER analyses relied on a single genome to represent each species, which conflates fixed changes separating species with polymorphic sites within a species. To overcome this limitation, we utilized T2T assemblies that have become available for diverse human and great ape individuals^32,35–38^. In this work, we leveraged these resources to identify 1,596 Consensus HAQERs, regions of rapid sequence evolution between the human-chimpanzee ancestor and an inferred human ancestral node, disentangling intraspecific polymorphism from lineage-specific divergence.

Consensus HAQERs show signatures consistent with elevated mutation rates, ancient positive selection, and bivalent regulatory function, reinforcing observations from our previous HAQER analysis^27^. Using single-cell, multiplexed enhancer assays in developing brain and heart cell types, we demonstrate that HAQERs can act as enhancers in multiple tissues, suggesting active roles in shaping human-specific gene regulation across multiple tissues and cell types.

## Results

### A 20-way telomere-to-telomere alignment of great ape divergence and variation

We leveraged newly available T2T genome assemblies for both great apes^32^ and diverse humans^35–38^ to construct a 20-way alignment that captures divergence between the great ape species and incorporates the genetic variation within each species. This alignment is referenced on the hs1/T2T-CHM13 human genome^33^, which is a sample with primarily European ancestry. To distinguish interspecies sequence differences between humans and the other great apes from polymorphic sites among human individuals, we included 7 additional diverse T2T human haplotypes: 2 from HG002 (maternal and paternal; Ashkenazi ancestry) and 5 from Han Chinese individuals (CN1, maternal and paternal; YAO, maternal and paternal; and Han 1)^35–38^. Beyond humans, we incorporated 12 T2T genome assemblies representing non-human apes, including primary and alternate haplotypes from each of chimpanzee, bonobo, gorilla, Bornean orangutan, Sumatran orangutan, and Siamang gibbon (**Figure 1A**). This alignment expanded upon the 9 haplotypes we previously analyzed^32^ with 11 new assemblies: 7 human haplotypes, 2 gibbon haplotypes, and 2 Bornean orangutan haplotypes (**Figure 1B**).

**Figure 1:**
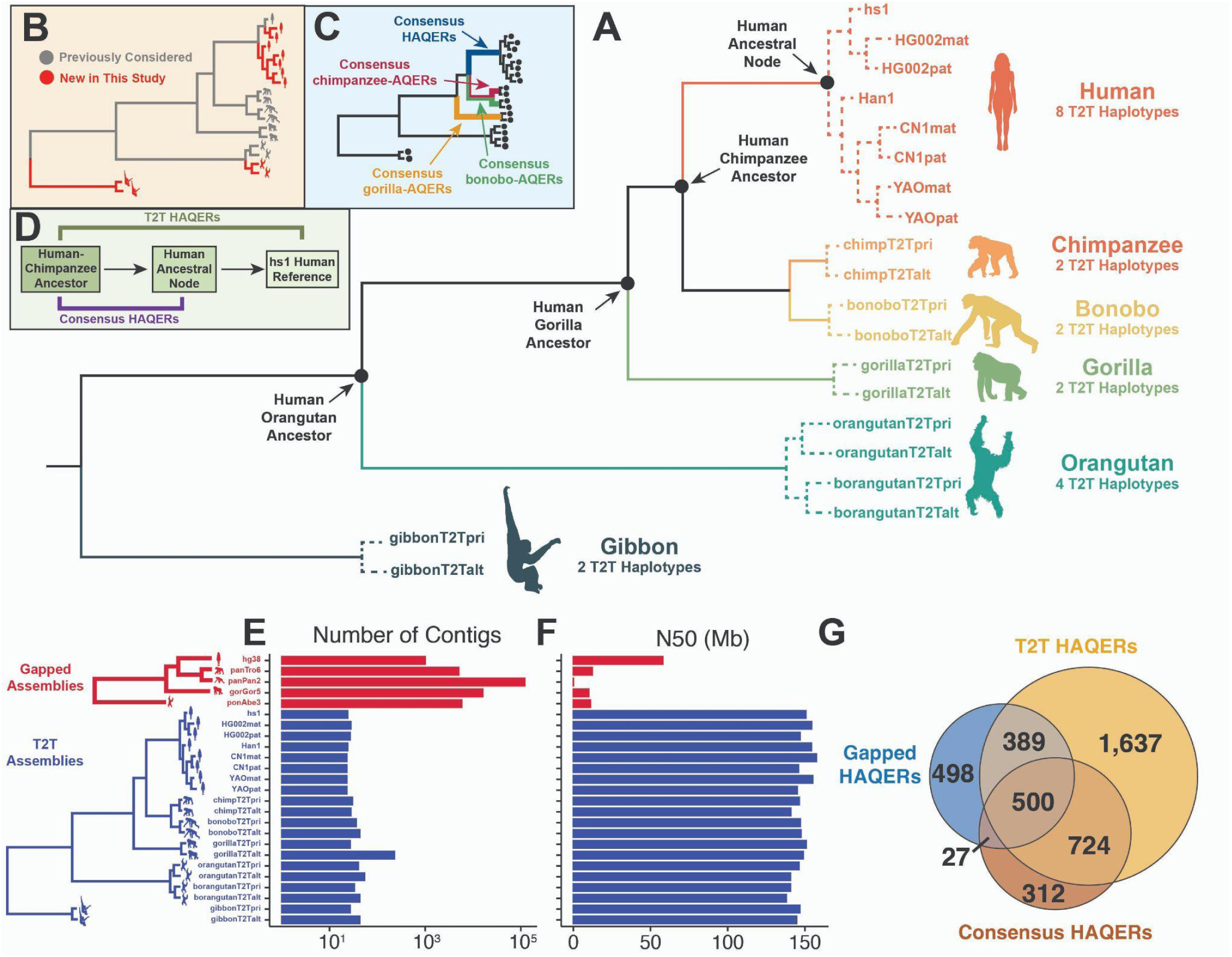
A 20-way alignment of telomere-to-telomere human and great ape genomes refined identification of rapidly evolved regions. (A) Phylogeny of a genome-wide 20-way alignment of haplotype-resolved telomere-to-telomere (T2T) assemblies spanning human and ape evolution and diversity. Key ancestral nodes are highlighted, as well as the number of assemblies per species. Dashed branches indicate intraspecific relationships. (B) Comparison of the 20-way alignment of T2T assemblies reported here with the previous 9-way alignment of T2T assemblies used to assess rapid divergence in Yoo et al^32^. (C) Branches of interest for ancestor quickly evolved regions (AQER) discovery. We define consensus human-AQERs (HAQERs) as regions of rapid evolution between the human-chimpanzee ancestor and the human ancestral node. Analogous AQER sets are identified for the chimpanzee, bonobo, and gorilla lineages. (D) Schematic contrasting the definition of T2T HAQERs^32^ with Consensus HAQERs (this study). T2T HAQERs measure divergence between the human-chimpanzee ancestor and the modern human assembly, hs1, whereas Consensus HAQERs measure divergence between the human-chimpanzee ancestor and the human ancestral node. (E-F) Assembly statistics comparing T2T (blue) and gapped (red) reference assemblies, including (E) the number of contigs and (F) contig N50 values. T2T genomes uniformly exhibit higher contiguity relative to gapped assemblies. (G) Number of overlaps among HAQER sets, including Gapped HAQERs^27^ (1,414 elements lifted to hs1 from 1,581 elements on hg38), 3,268 T2T HAQERs^32^, and 1,596 Consensus HAQERs (this study). Total set size may not equal the reported overlap sizes as individual regions can overlap multiple regions in another set. While, at the high level, we would expect every Consensus HAQER to be a Gapped HAQER and T2T HAQER, many are not Gapped HAQERs due to the missing genomic fragments in those assemblies, and not T2T HAQERs due to alignment changes between human segments and non-human segments, when adding an additional 7 human haplotypes to the alignment.

These T2T assemblies exhibit dramatically higher contiguity relative to pre-T2T assemblies, which we refer to as “gapped” assemblies, as measured by both number of contigs and contig N50 values (**Figure 1E-F**). Critically, T2T assemblies introduced hundreds of megabases missing from gapped assemblies, enabling the identification of evolutionary innovations in previously unexplored regions of the genome.

### Incorporating intraspecific variation into the identification of rapidly evolved genomic regions

In previous work, we developed a computational screen to identify the most divergent regions of the human genome, which we called HAQERs, for Human Ancestor Quickly Evolved Regions^27,32^. Our first implementation began with a 5-way genome-wide alignment of humans and great apes, using pre-T2T (gapped) reference assemblies (human hg38, chimpanzee panTro6, bonobo panPan2, gorilla gorGor5, and Sumatran orangutan ponAbe3). Based on this genome-wide multiple alignment and the standard species topology, we inferred the human-chimpanzee ancestral sequence. We then identified HAQERs as regions with significantly elevated molecular divergence (inclusive of substitutions, short insertions, and short deletions) between this inferred human-chimpanzee ancestor and the modern human reference genome (at least 29 mutations in 500bp). This screen produced a set of 1,581 elements we refer to as *Gapped HAQERs*^27^.

In the second iteration of HAQER identification, we applied this same framework to a 9-way alignment of T2T genomes, which allowed us to search regions of the human genome that had not been present in the gapped human reference assembly, or where there were not a sufficient number of other great ape assemblies that contained the orthologous segment. This produced an expanded set of 3,268 *T2T HAQERs*^32^.

Notably, T2T HAQERs are not merely a superset of Gapped HAQERs, as 525 regions in the Gapped HAQER set do not overlap the T2T HAQER set. We reasoned that this discrepancy could be explained by the fact that the T2T human reference genome is not strictly the gapped reference sequence with the gapped regions included; rather, the T2T genome is derived from a different human sample of distinct continental ancestry. Thus, some divergence signals may change due to variation between human individuals that arose after the human coalescent rather than fixed differences between lineages^33,39^. Similarly, gapped and T2T non-human primate assemblies were also derived from distinct ape individuals^32,40^.

Prompted by this observation, in this study, we define *Consensus HAQERs* as the most divergent regions separating the inferred human-chimpanzee ancestor from an inferred human ancestral node, the most recent common ancestor of all extant human haplotypes in our 20-way alignment (**Figure 1D**). We identified 1,596 Consensus HAQERs, covering approximately 1.7 million bases and 0.05% of the human genome.

Our current study leverages probabilistic base representations^41^ to a greater degree than our previous work. To represent ancestral sequence, we assign a vector of probabilities encompassing A, C, G, and T to each ancestral base that is inferred to be present at an alignment position. Previously, we measured divergence as the number of sequence differences between the reconstruction of the human-chimpanzee ancestor and a modern human genome. Our current study measures divergence between the human-chimpanzee ancestor and a human ancestral node, both internal nodes with vectors of probabilities, as one minus the cosine distance between their probabilistic base representations. This captures the likelihood of a mutation having occurred on the branch separating the two internal nodes (see **Methods**).

Beyond humans, we applied this same framework to identify rapid evolution across other lineages, identifying 2,517 chimpanzee-AQERs, 2,403 bonobo-AQERs, and 523 gorilla-AQERs (**Figure 1C**). Similar to humans, each of these sets was defined from a 20-way T2T alignment referenced on the primary extant haplotype between the common ancestor of the extant haplotypes and the common ancestor between that species and humans (**Figure 1C**). We defined a uniform statistical threshold for AQER identification for all lineages (pAdj < 3e-7 using the maximum divergence rate in a 10 Mbp window as a conservative estimate of the expected divergence rate; see **Methods**), which in a 500 bp window corresponds to 29 mutations for all 3 HAQER sets, chimpanzee-AQERs, and bonobo-AQERs, and 36 mutations for gorilla-AQERs. The relatively small number of gorilla-AQERs resulted from the longer phylogenetic branch between the human-gorilla ancestor and extant gorillas in the standard phylogeny, which increased the branch length-adjusted divergence threshold and made detection more conservative. In all cases, consensus AQER sets are smaller than the corresponding T2T sets, consistent with the fact that consensus AQERs extend only to the species coalescent point rather than to mutations that accumulated along branches leading to modern sampled individuals.

When comparing HAQER sets, we observe considerable overlap, but also subsets specific to one of the analyses. Regions specific to the Gapped HAQER set are largely attributable to within-human polymorphic alleles of hg38 erroneously counted as divergence between the human-chimpanzee ancestor and the modern human (**Figure 1G**). Those regions specific to T2T HAQERs are the result of the intraspecific polymorphic alleles of hs1 being conflated with interspecies divergence in a larger screen spanning more of the genome. In this work, we proceed to focus on Consensus HAQERs, which are based on a genome-wide alignment of diverse T2T assemblies, making it possible to control for human variation when identifying the most rapidly diverged segments in the human genome.

### HAQERs exhibit elevated mutation rates

We next sought to uncover the forces underlying the dramatic divergence in HAQER regions. We hypothesized that heightened divergence could be due to a longer coalescent time, an elevated mutation rate, or selective pressure.

Because our approach uses a fixed-topology phylogenetic model, we evaluated whether incomplete lineage sorting (ILS)^42^, where some parts of the genome do not follow the species tree, could contribute to elevated interspecies divergence. Overlaps between Consensus HAQERs and inferred genome-wide ILS states^43^ (**Figure S2D**) revealed that HAQERs are significantly depleted from the nonstandard human-gorilla ILS topology, indicating that the elevated human divergence of the Consensus HAQER set is unlikely to be the result of human-gorilla ILS. Instead, HAQERs are enriched in regions annotated as the canonical great ape topology, but with extended human branch lengths, consistent with accelerated substitutions on the human lineage. We also observed a modest enrichment in chimpanzee-gorilla ILS regions, which can mimic patterns of human-specific rapid divergence as both scenarios yield excess divergence between humans and other apes. These intersections suggest that ILS may contribute to some individual HAQER elements, but is not a sufficient explanation for their rapid divergence in the human genome.

One approach to understand if elevated mutation rates are contributing to dramatic divergence is analyzing sequence features associated with elevated local mutation rates. We first examined whether HAQERs are preferentially identified within repetitive elements^44,45^. All 3 HAQER sets are significantly depleted from the general repeat classes of LINEs (Long Interspersed Nuclear Elements) and SINEs (Short Interspersed Nuclear Elements), yet significantly enriched for particular families of these elements, such as SVAs (SINE-VNTR-Alu) (Gapped HAQERs: pAdj < 1e-29, T2T HAQERs: pAdj < 1e-17, Consensus HAQERs: pAdj < 1e-13). This enrichment in SVA elements is due to rapid divergence of the VNTR region within the SVA, and VNTRs have long been associated with a high mutation rate^46,47^. We also observe other enrichments for repeat classes that have been associated with high mutation rates, such as simple repeats^48,49^ and segmental duplications^45^ (**Figure 2A**). Notably, satellites are depleted in Gapped HAQERs but enriched in T2T– and Consensus HAQERs, consistent with the improved assembly of satellite DNA in T2T genomes. We identified similar repeat enrichment patterns in non-human primate AQERs (**Figure S1**), suggesting that many of these features may not be human-specific, but instead general features of primate genome evolution. By contrast, simple repeats, which are known to be associated with increased mutation rates, are uniquely enriched in HAQERs and depleted in ape AQERs. SVA element enrichments are also specific to HAQERs and not observed in ape AQERs, consistent with the recent origin of SVAs and their increasing internal VNTR length in the human lineage^50,51^. Overall, HAQERs are enriched for repeat classes that are associated with increased mutation rates.

**Figure 2:**
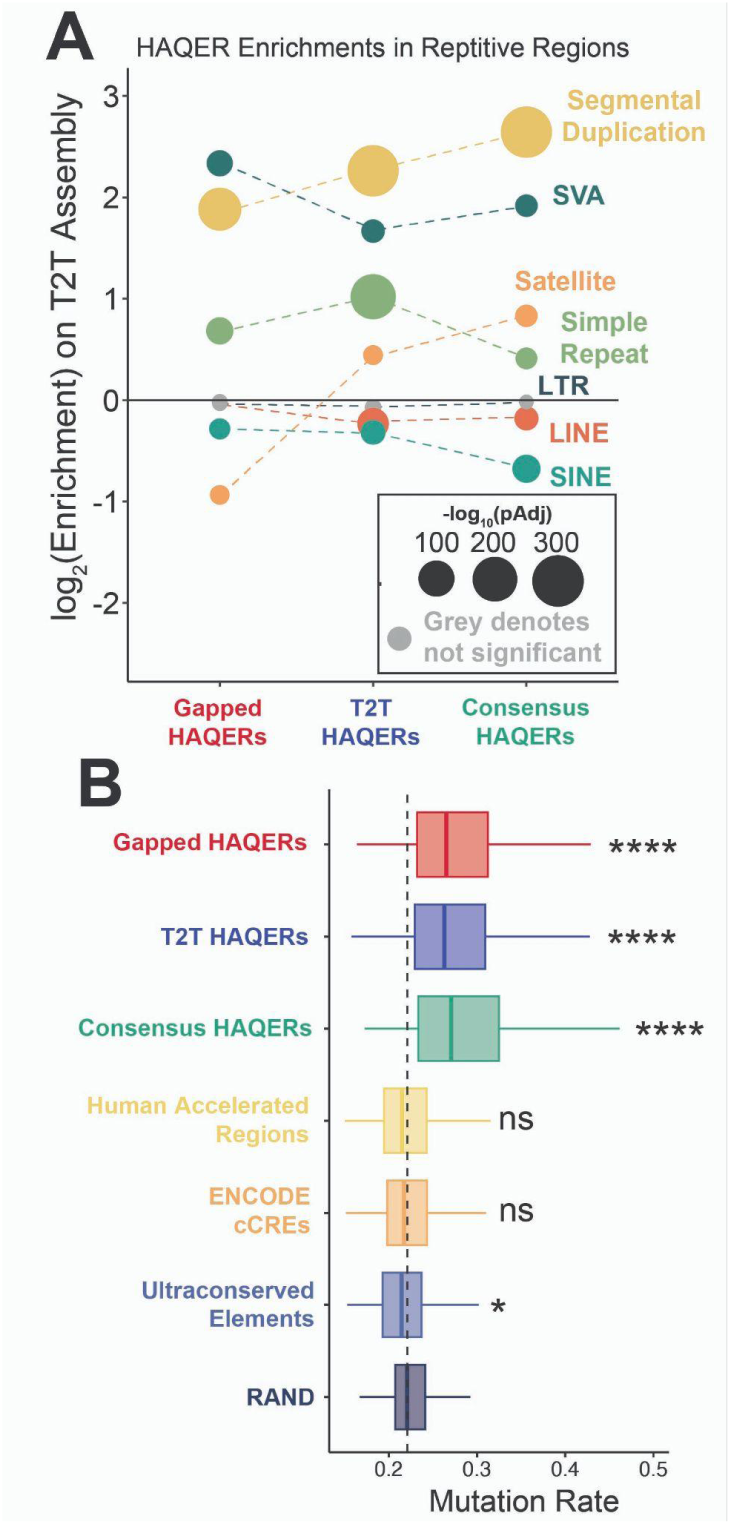
HAQERs exhibit elevated mutation rates. (A) Enrichment of Gapped HAQERs, T2T HAQERs, and Consensus HAQERs for overlapping repetitive element classes and families, shown as log2 fold enrichment relative to a uniform distribution over the genome. Repeat classes and families that have been associated with elevated mutation rates, such as segmental duplications, SVAs (which contain VNTRs), and simple repeats, show an enrichment for overlapping HAQER sets. Point size indicates statistical significance, with nonsignificant enrichments (pAdj > 0.05) in grey. (B) Mutation rate estimates across regions of evolutionary interest, including HAQER sets, human accelerated regions (HARs), ENCODE candidate *cis*-regulatory elements (cCREs), ultraconserved elements, and random neutral proxy genomic regions (RAND). (Bonferroni-corrected pairwise t-test vs. RAND; * pAdj < 0.05; **** pAdj < 0.0001; ns = not significant).

We next examined the GC content of HAQERs, given known associations between GC content and mutation rates^44,52^. Compared to the genome-wide distribution, HAQERs are enriched for both GC-rich and GC-poor regions (**Figure S2A-C**). GC-rich regions have been associated with higher mutation rates, in part due to the elevated rate of CpG to TpG changes^53,54^, and GC-poor HAQERs may also contribute to an overall greater rate of mutation through repeats^55^.

To more directly assess mutation rates, we intersected HAQERs with base-level estimates derived from local patterns of *de novo* mutations^56^ (**Figure 2B**). HAQER sets display pronounced and significantly higher mean mutation rates, supporting elevated local mutagenesis as a mechanism driving rapid HAQER evolution (Bonferroni-corrected pairwise t-test against random genomic regions (RAND); Gapped HAQERs: pAdj < 1e-50, T2T HAQERs: pAdj < 1e-50, Consensus HAQERs: pAdj < 1e-50). By contrast, mutation rates in human accelerated regions (HARs)^6^ and ENCODE candidate *cis*-regulatory elements (cCREs)^57^ are indistinguishable from the genomic background. Ultraconserved elements (UCEs)^58^ exhibit borderline significance (pAdj ≈ 0.01) for reduced mutation rates. Taken together, these findings suggest that elevated rates of mutagenesis supported HAQER evolution.

### HAQERs exhibit signatures of ancient positive selection

We investigated whether HAQER divergence reflects not only elevated mutation rates but also positive selection. A general signature of ancient positive selection is an excess of divergence relative to estimates of mutation rate.

Under neutral evolution, divergence and polymorphism tend to be positively correlated^59^. Positive selection, however, increases the fixation rate of beneficial variants, increasing the divergence-to-polymorphism ratio. Ancient beneficial variants, emerging millions of years ago, have had enough time to complete selective sweeps, become fixed in the population, and regain polymorphism through additional mutations. However, the polymorphism-to-divergence ratio will still be lower than expected due to positive selection^60,61^. Thus, we would expect ancient positive selection to lead to both a low proportion of divergent alleles that are polymorphic, and a low ratio of polymorphic alleles to divergent sites within a region. In general, a low ratio of the mutation rate estimate to the amount of divergence can be a signature of ancient positive selection.

Consistent with this signature, all three HAQER sets exhibit significantly higher proportions of fixed divergent sites, when compared to HARs and RAND (Chi-squared test against RAND; Gapped HAQERs: p < 1e-50, T2T HAQERs: p < 1e-50, Consensus HAQERs: p < 1e-50) (**Figure 3A**). HARs also exhibit an enrichment for fixed sites (p < 1e-13). The disproportionately high fixation of divergent alleles is consistent with many HAQERs carrying beneficial changes that were subjected to positive selection after the split from chimpanzees. By contrast, missense mutations demonstrate the opposite pattern (p < 1e-50), suggesting long-term negative selection, consistent with past reports^27^. These trends persist when we normalize the number of divergent sites to an estimation of the mutation rate that is the count of all polymorphic sites within each region set (**Figure 3B**). We then normalized sequence divergence in each set to estimates of local mutation rates^56^, finding that HAQERs exhibit an abnormally high ratio of divergent sites to mutation rate (**Figure 3C**). These complementary analyses revealed that the divergence in HAQERs exceeds estimates of their mutation rates and presents the possibility that many have disproportionately acquired fixed derived alleles, suggesting historic positive selection.

**Figure 3:**
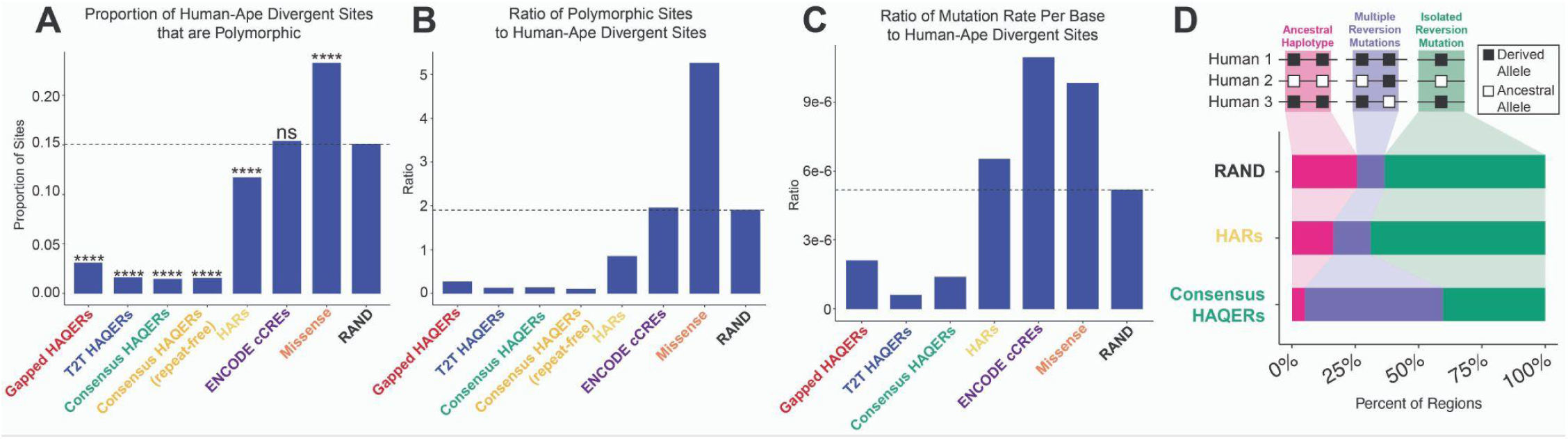
HAQERs show evidence of ancient positive selection. (A) Proportion of human-ape divergent sites that are polymorphic across region sets. HAQER sets exhibit the lowest fractions of polymorphic alleles. (Chi-squared vs. RAND; **** p < 0.0001; ns = not significant) (B) Ratio of all polymorphic sites (including non-divergent sites) to divergent sites across region sets. (C) Ratio of average local mutation rate to human-ape divergent sites across region sets. (D) Relative frequencies of potential reversion mutations and ancestral haplotypes among polymorphic divergent alleles with high derived allele frequencies (> 0.95).

While HAQERs were ascertained based on divergence, they clearly deviate from the genome-wide relationship between divergence and estimates of the mutation rate (**Figure S3B-C**). They exhibit rapid divergence without elevated polymorphisms, further evidence for non-neutral evolution driven by positive selection. A potential concern is that polymorphisms are more difficult to map in repetitive regions, where HAQERs are enriched, potentially deflating observed polymorphic rates. To mitigate this, we analyzed a repeat-free subset of Consensus HAQERs, excluding HAQERs overlapping simple repeats, satellites, segmental duplications, or mobile elements. Even in this conservative set, the proportion of polymorphic divergent alleles remained significantly lower than RAND (**Figure 3A**), indicating that this signal is not attributable to mapping artifacts.

All HAQER sets show an excess of high frequency derived alleles (**Figure S3G**), which we hypothesized could be due to incomplete fixation and recent positive selection, or reversion mutations following ancient fixation. To distinguish between these possibilities, we examined the local sequence context around polymorphic divergent sites with high derived allele frequencies (> 0.95), assessing whether ancestral alleles occur on a haplotype with other ancestral alleles (consistent with incomplete sweeps), or without any nearby ancient alleles on the haplotype (greater likelihood of reversion). Notably, HAQERs exhibit a considerably higher proportion of potential reversion mutations compared to HARs and RAND, consistent with ancient positive selection in regions with elevated mutation rates that cause reversion mutations (**Figure 3D**).

Because ancient positive selection fixes many variants, we would expect to detect negative selection acting on these sites in the present day. We observed inconsistent signatures of recent negative selection (**Figure S3**), with substitution variant calls on hg38 showing an excess of rare alleles in HAQER sets, but we do not observe this same excess when using variant calls mapped to the hs1 assembly. Previous work has reported that HAQERs are enriched for rare structural variant alleles, which would be consistent with negative selection in modern humans^29^.

Taken together, HAQERs exhibit signatures consistent with ancient positive selection and mixed signatures of negative selection in the present day.

### HAQERs are enriched in bivalent chromatin states

Having observed signatures that HAQERs evolved through elevated mutation rates and positive selection, we next investigated the associations of these elements with gene regulation. To this end, we first analyzed overlap enrichments between HAQER sets and chromatin state annotations across 127 reference epigenomes^62^, spanning diverse epigenomic contexts and body systems. All HAQER sets are enriched in bivalent chromatin states (both enhancers and promoters), polycomb repressed regions, and ZNF/repeat states, while depleted from active enhancers and transcribed regions (**Figure 4A**). Bivalent chromatin states carry both activating (H3K4me3/H3K4me1) and repressive (H3K27me3) histone marks, leaving promoters and enhancers reversibly silenced but poised for context-specific activation^63–65^. T2T HAQERs show weaker enrichments than Gapped HAQERs, likely due to the added discovery of HAQERs in non-regulatory regions, such as near centromeres. Consensus HAQERs exceed T2T enrichments, supporting their functional relevance and a potential tendency for rapid evolution to occur in context-specific regulatory elements^30^.

**Figure 4:**
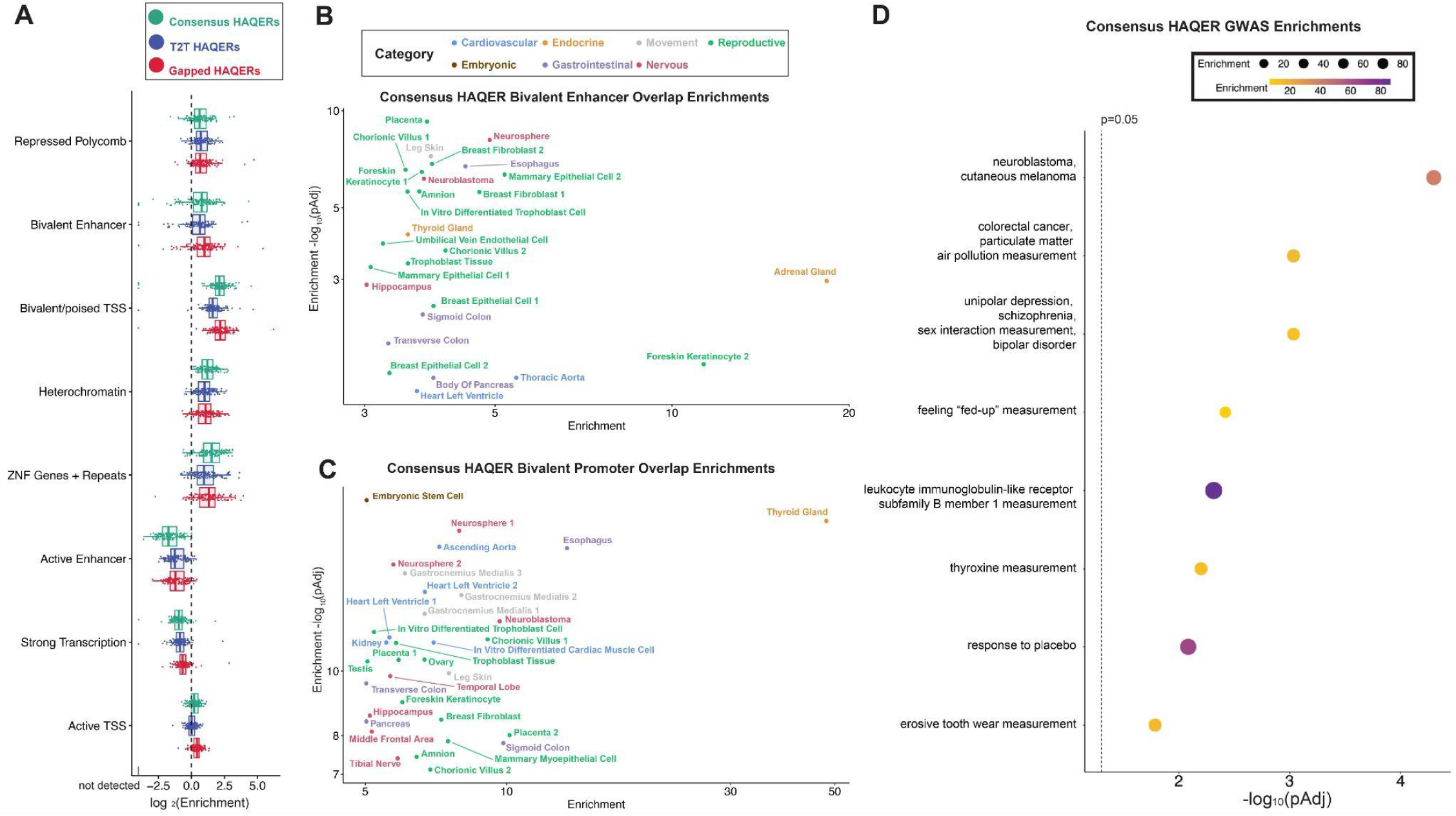
HAQERs are enriched in bivalent chromatin states and disease-linked loci. (A) Chromatin state enrichments for Consensus, T2T, and Gapped HAQERs across 127 reference epigenomes^62^. HAQERs are enriched for bivalent states, polycomb repression, and ZNF/repeats, while depleted from active enhancers and transcribed regions. (B-C) Volcano plot of Consensus HAQER enrichments in (D) bivalent enhancer and (E) bivalent promoter chromatin states across tissues and cell types. (D) FDR-corrected p values for overlap enrichments between Consensus HAQERs and GWAS-linked loci. Dotted line marks p = 0.05.

We next examined Consensus HAQER enrichments in bivalent states across an expanded set of 833 epigenomes to better understand the specific tissues and cell types affected.^17^ In total, 125 Consensus HAQERs overlap bivalent enhancers and 70 overlap bivalent promoters, with many Consensus HAQERs annotated as bivalently poised across numerous tissues and cell types (**Figure S4A-B**). We observed the strongest enrichments across several organ systems, including nervous, reproductive, cardiovascular, and gastrointestinal tissues (**Figure 4B-C**), consistent with prior studies of human-specific development and gene regulation that noted significant changes in these tissues^1,66–69^. Similar bivalent enrichments persist in HAQER subsets defined by stricter divergence thresholds or the exclusion of repetitive elements (**Figure S4C**).

The window size used when calculating the acceleration or velocity of human genetic divergence has varied across different studies. Early screens of acceleration used smaller window sizes^18,19^ than the HAQER screens for velocity over 500 bp^27,32^. We hypothesized that the genomic scale used by the screen would influence the function of the highly divergent regions identified. We therefore repeated HAQER identification across a range of sliding window sizes from 25 bp to 20 kbp to capture rapidly evolved elements across multiple spatial scales (**Figure S4D**). Some HAQERs are only detectable at specific window sizes, and HAQERs at larger spatial scales are often composites of multiple HAQERs identified at smaller spatial scales. Bivalent state enrichments fluctuate across window sizes, peaking at around 1 kbp for enhancers and 200 bp for promoters, consistent with the typical sizes of these regulatory elements^70,71^ (**Figure S4E**). This suggests that intermediate window sizes (200 bp to 1 kbp) may be the unit sizes of regulatory evolution where sequence divergence is most correlated with gene regulatory divergence. Very small windows of divergence may be dominated by the stochastic grouping of mutations^72^, and very wide windows may capture large-scale fluctuations in mutation rate along the chromosome^73^.

### HAQERs are enriched in disease-linked loci

To assess potential phenotypic impacts associated with the divergence in Consensus HAQERs, we quantified enrichments between Consensus HAQERs and genome-wide association study (GWAS) loci from the GWAS Catalog^74^ (see **Methods**). There is a consistency between the tissues and cell types most enriched for bivalent regulatory elements, and the categories of GWAS loci that tend to overlap HAQERs. Neuroblastoma has some of the highest enrichments for both bivalent enhancer and bivalent promoter chromatin states overlapping Consensus HAQERs, and Consensus HAQERs are also enriched for GWAS SNPs (single nucleotide polymorphisms) associated with neuroblastoma (**Figure 4B-D**). More generally, we see strong enrichments for bivalent states related to brain development (e.g. neurosphere) and GWAS loci related to psychiatric disorders. A second pair of matching enrichments is the tendency of Consensus HAQERs to overlap both bivalent regulatory elements from samples related to the gastrointestinal tract and GWAS loci related to colorectal cancer (**Figure 4B-D**). A third pair of matching enrichments is between bivalent chromatin states in the thyroid gland and GWAS loci associated with thyroxine levels (**Figure 4B-D**). While we have primarily focused on Consensus HAQERs, there is also a complementary set of ‘variable’ HAQERs that are not repeatedly identified as significantly divergent across human haplotypes, composed of Gapped HAQERs not overlapping T2T HAQERs and T2T HAQERs not overlapping Gapped HAQERs. This set shows a large number of enrichments, partially due to its large set size. The enrichments span psychiatric and neurodegenerative disorders, as well as immune, metabolic, and cancer-related traits (**Figure S5**). Overall, these results are consistent with divergence in Consensus HAQERs having shaped human phenotypes related to the nervous, gastrointestinal, and endocrine systems. Not only has this divergence shaped all humans, but present-day variation among humans within these highly-divergent regions continues to influence phenotypes related to our brain, gut, and hormones.

### Multiple HAQERs are active enhancers in neurodevelopment

Based on HAQER enrichments for both regulatory chromatin states in neurodevelopmental cell types and neuropsychiatric risk loci, we assayed 8 HAQERs for gene regulatory function in cell types from the developing mouse brain to confirm their function. We prioritized candidate HAQERs overlapping open chromatin peaks in the developing brain^62^. The set included 7 Consensus HAQERs and 1 variable HAQER (Gapped HAQER1463). We assayed enhancer activity in the developing mouse brain using *in vivo* scSTARR-seq (single-cell self transcribing active regulatory region sequencing)^27^. HAQER sequences were cloned into a plasmid library and co-injected with a constitutive GFP (green fluorescent protein) reporter into embryonic mouse cerebral cortices via *in utero* electroporation at E14.5. In contrast to our initial study describing scSTARR-seq, where cells were harvested ∼18 hours after injection, we recovered cells after 2 days to obtain more cells and a broader representation of neuronal types. Following dissection, we used FACS (fluorescence-activated cell sorting) to isolate GFP+ transfected cells for single-cell RNA sequencing. Each isolated cell represents a multiplex enhancer assay experiment, and the cells encompass diverse developmental cell types (**Figure 5A**).

**Figure 5:**
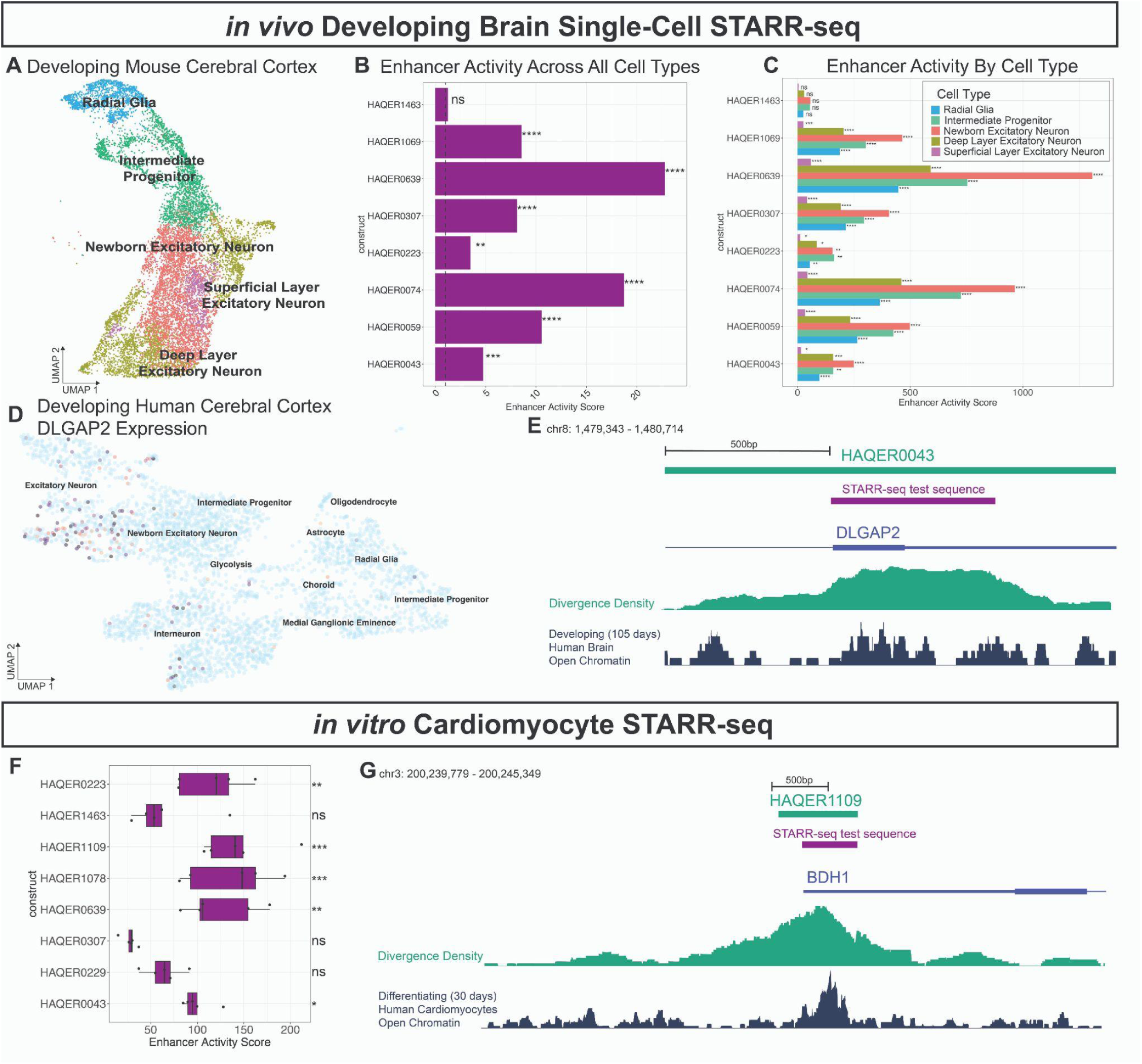
HAQERs function as active enhancers in the developing brain and developing human heart. (A) UMAP (uniform manifold approximation and projection) of 12,435 single cells collected from embryonic mouse cortices 2 days after *in utero* electroporation at E14.5 with HAQER scSTARR-seq (self transcribing active regulatory region sequencing) libraries. (B) Pseudobulk enhancer activity for each construct, quantified as reporter expression relative to a GFP transfection control. Seven of eight HAQERs show significant activity compared to random scrambled negative control sequences. (C) Cell-type-resolved enhancer activity across excitatory developmental lineages, with robust activity for 7 Consensus HAQERs. (D) Expression of *DLGAP2* in a single-cell transcriptional profile of the developing human cortex, compiled from samples across stages of peak neurogenesis^75^. (E) HAQER0043, an active enhancer in the developing cerebral cortex, overlaps an exon of *DLGAP2*, a synaptic scaffold gene linked to neuropsychiatric and neurodegenerative disorders^76^. (F) Pseudobulk enhancer activity of 8 HAQERs assayed in iPSC-derived cardiomyocytes. (G) HAQER1109, a cardiomyocyte enhancer, overlaps the 3’ UTR of *BDH1*, a ketone metabolism gene associated with oxidative stress during heart failure^77^. (* p < 0.05; ** p < 0.01; *** p < 0.001; **** p < 0.0001; ns = not significant)

We first quantified pseudobulk enhancer activity for each HAQER construct. We calculated the enhancer activity for each HAQER as the STARR-seq reporter RNA to DNA ratio, relative to the same ratio for the random scrambled negative control constructs, pooling reads from all cells (see **Methods**). We found that all Consensus HAQERs (but not Gapped HAQER1463) exhibited significant enhancer activity in the pseudobulk analysis (**Figure 5B**). We then leveraged the single-cell expression profiles to deconvolve enhancer activity into the activity of the individual cell types. The Consensus HAQERs demonstrated significant enhancer activity across all 5 cell types of the excitatory developmental lineage that we were able to identify (**Figure 5C**). Among the tested HAQERs, HAQER0639 demonstrated the highest enhancer activity in both pseudobulk and cell type-specific analyses. Notably, HAQER0043 overlaps an intron-exon boundary in *DLGAP2*, a gene encoding a postsynaptic scaffold protein linked to schizophrenia, autism spectrum disorder, and Alzheimer’s disease^76,78^ (**Figure 5E**). Consistent with the enhancer activity patterns of HAQER0043, *DLGAP2* is expressed in excitatory neurons, particularly newborn neurons, in the developing human cerebral cortex^75^ (**Figure 5D**). Together, these data identify multiple HAQERs as active enhancers in the developing cerebral cortex, which may coordinate human-specific transcriptional programs and disease susceptibility.

### HAQERs as active enhancers in human heart development

Beyond the brain, we observed HAQER enrichment in bivalent chromatin states in multiple heart-related reference epigenomes (**Figure 4D-E**). Based on this observation, we expanded scSTARR-seq to human iPSC (induced pluripotent stem cell)-derived cardiomyocytes to understand if HAQERs function as enhancers in the developing human heart (**Figure S6**). We prioritized 8 candidates (6 Consensus HAQERs and 2 variable HAQERs: HAQER0229 and HAQER1463) based on regulatory chromatin state overlaps in heart-related epigenomes. Five of these regions also overlap open chromatin elements in the developing brain, and were included in both STARR-seq assays. We transfected the STARR-seq plasmid library into iPSC-derived human cardiomyocytes and after 2 days sequenced 5,648 cells. Five of these eight HAQERs exhibited significant enhancer activity (**Figure 5F**). HAQER1109 exhibited robust enhancer activity in cardiomyocytes, and overlaps the 3’ UTR (untranslated region) of *BDH1*, a gene encoding a ketone metabolism enzyme, whose overexpression in heart tissue ameliorates oxidative stress during heart failure (**Figure 5G**)^77^. These results extend scSTARR-seq to human heart cells and highlight HAQERs as candidate cardiomyocyte enhancers with disease relevance.

## Discussion

The emergence of T2T genome assemblies for both humans^33,35^ and great apes^32^ offers opportunities to explore genome evolution in previously inaccessible genomic regions, including repetitive regions such as centromeres, telomeres, and segmental duplications. We anticipate that these historically underexplored regions harbor underrecognized functional elements relevant to human evolution and disease. As such, detailed comparative analyses are required to uncover the evolutionary significance of these new regions. Here we begin to use these new T2T resources, constructing a 20-way alignment of near-complete genome assemblies spanning human and great ape diversity to define Consensus HAQERs, which represent the 1,596 most divergent regions between the human-chimpanzee ancestor and an ancestral node of modern humans. Much of comparative genomics has historically relied on comparisons between single reference genomes, which are unable to distinguish fixed sites separating lineages from polymorphic alleles within populations. We find substantial disagreement between HAQER sets ascertained on different sets of reference genomes, such as Gapped HAQERs (ascertained on pre-T2T gapped genome assemblies) and T2T HAQERs (ascertained on T2T genome assemblies), establishing the importance of capturing intraspecific variation when identifying regions of interspecific divergence. Our current approach, informed by probabilistic ancestral reconstruction incorporating intraspecific variation, mitigates this reference bias. We leveraged available T2T human assemblies, including one European haplotype, two Ashkenazi haplotypes, and five Han Chinese haplotypes. Notably, this set lacks T2T genomes of African ancestry, which are needed to represent a major component of human genetic diversity. The future availability of more diverse T2T genomes will be valuable to further refine the set of regions that consistently show elevated divergence across all humans.

Consistent with prior findings^27,79,80^, HAQERs exhibit evidence of both elevated mutation rates and ancient positive selection. Local mutation rates are highly variable across the genome^56^. Recent studies have clarified that mutation rate nonuniformity is linked to genomic features, with open chromatin elements exhibiting higher mutation rates^81^ and gene bodies exhibiting lower mutation rates^82^. Notably, nearly half of known mutations underlying adaptive traits in modern humans occur in mutation-prone regions^29,79,83^. These observations emphasize that regions of elevated mutation rates should be recognized as key substrates of adaptation.

While our analyses revealed signatures of ancient positive selection in HAQERs, these analyses were conducted at the level of HAQER sets as a whole. We expect that selection in these regions is heterogeneous, with the divergence in some HAQERs being under selection and other HAQERs evolving neutrally. Developing robust methods to evaluate selection at single loci across deep evolutionary timescales remains an important future challenge.

HAQERs are preferentially enriched in bivalent chromatin states, with this current dataset now reinforcing our previous finding^27^ that lineage-specific sequence divergence preferentially targets regulatory elements with context-specific roles in development and environmental response^63–65^. This enrichment is strongest in subsets defined by higher divergence thresholds, suggesting that the fastest-evolved regions of the genome are disproportionately regulatory. HAQERs also tend to occur in GC-rich regions, consistent with promoter and enhancer activity. Further analysis of sequence divergence across spatial scales revealed that the length over which divergence is calculated influences the functions impacted by that divergence.

To more directly test the functional roles of HAQERs, we performed multiplex single-cell enhancer screens to verify that HAQERs act as enhancers in both the developing brain and heart. These include a HAQER acting as an intronic brain enhancer in the *DLGAP2* locus, a gene implicated in autism spectrum disorder and schizophrenia, and a HAQER within the 3’ UTR of BDH1 that has enhancer activity in cardiomyocytes. Also in support of Consensus HAQERs influencing human biology are enrichments for diverse GWAS loci, including cancer, neuropsychiatric disorders, and endocrine traits. Together, these results establish Consensus HAQERs as a valuable catalog of rapidly evolved regions separating humans from other great apes that we anticipate to contain human-altered gene regulatory elements and underlie human-specific traits and disease susceptibilities.

### Limitations of the study

There are limitations due to uncertainties in the whole-genome alignments upon which AQERs were identified. While T2T genomes have made highly repetitive regions available to analyze, many of these regions, particularly centromeres and telomeres, are so rapidly evolving that we were not able to effectively generate alignments across them, which means that they were not included in our analysis. When additional human T2T assemblies were added to the genome-wide multiple alignment, the positioning of non-human orthologs changed, resulting in Consensus HAQER regions that do not overlap T2T HAQERs. For example, in one genomic location, non-hs1 human haplotypes have an insertion of 20 to 24 bp that caused ape orthologs to adjust their alignment in the area and lead to a higher estimated number of mutations on the branch to humans.

While we have demonstrated the functional relevance of HAQERs in gene regulation, future work will be required to demonstrate how HAQER evolution has altered gene expression and organismal phenotypes.

We detected relatively few gorilla-AQERs, likely due to the long phylogenetic branch from the human-gorilla ancestor, indicating that our framework for identifying functionally significant mutations across lineages may be best tuned for closely related species.

An important open question is the relative contribution of distinct classes of genomic changes in human evolution. Human accelerated regions (HARs), which show rapid divergence in previously highly conserved elements^6,19^, likely reflect modifications of existing functional elements. This is also true for human conserved deletions (hCONDELs), which modified existing functions by eliminating them^84,85^. In contrast, HAQERs are identified as highly divergent regions without requiring prior conservation, allowing many to represent *de novo* regulatory elements in the human lineage. The balance among these mechanisms (modification of conserved elements versus *de novo* function) remains unclear^26^.

## Acknowledgments

We thank the Duke Cancer Institute Flow Cytometry Core, as well as the Duke Human Vaccine Institute Research Flow Cytometry Core and Viral Genome Analysis core facilities. We thank Luke Bartelt, Anushka Katikaneni, Hailey Napier, and Natalie Dzikowski for critical feedback. This research was supported by the Duke Whitehead Scholarship, and theNational Human Genome Research Institute (R35HG011332), the National Institute of General Medical Sciences (T32 GM007748), the National Institute of Neurological Disorders and Stroke (R37NS11038), the National Institute of Mental Health (R01MH132089), and the National Heart, Lung, and Blood Institute (R01HL157277) of the National Institutes of Health.

## Author Contributions

Y.L., R.J.M., and C.B.L. designed the study and wrote the paper. Y.L., R.J.M., S.W., S.A.Z., F.M., and M.T. performed experiments and analyses. M.K., R.K., D.L.S., and C.B.L. supervised and funded research. All authors reviewed, edited, and approved the manuscript prior to submission.

## Resource Availability

### Lead contact

Further information and requests for resources and reagents should be directed to and will be fulfilled by the lead contact, Craig B. Lowe.

### Materials availability

This study did not generate new unique reagents.

### Data and code availability

● Supplementary Tables can be accessed at 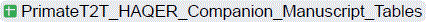
● Many software programs written for this manuscript were implemented as a part of Gonomics, an ongoing effort to develop an open-source genomics platform in the Go programming language (golang). Gonomics can be accessed at https://github.com/vertgenlab/gonomics.
● Additional software, as well as raw and analyzed datasets, including browser tracks, sequencing files, multiple alignments, variant sets used in selection analysis, and figure generation pipelines, will be made available at the time of publication on our lab website at https://www.vertgenlab.org. Additional raw and analyzed datasets will be deposited at GEO, to be made publicly available as of the date of publication.
● Any additional information required to reanalyze the data reported in this paper is available from the Lead Contact upon request.

### Experimental model and subject details

● Wild type CD1 mouse embryos at stage E14.5 were used for *in utero* electroporations as described in method details. We did not restrict our scSTARR-seq assays to embryos of only one sex; our data includes both developing males and females.
● All experiments were performed in agreement with the guidelines from the Division of Laboratory Animal Resources from Duke University School of Medicine and the Institutional Animal Care and Use Committee of Duke University.

## Method Details

### Genome assembly statistics

We calculated the number of contigs and the N50 of each genome assembly with gonomics: assemblyStats^86^.

### AQER set identification

#### Genome-wide multiple alignment

To construct a 20-way alignment spanning human and great ape divergence and diversity, we accessed 20 genome assemblies listed in the following table.

| ID | Description | Source |
| --- | --- | --- |
| hs1 | T2T reference human | https://hgdownload.soe.ucsc.edu/goldenPath/hs1/bigZips/hs1.fa.gz |
| HG002 | T2T non-reference human HG002 | https://s3-us-west-2.amazonaws.com/human-pangenomics/T2T/HG002/assemblies/hg002v1.0.1.fasta.gz |
| YAOmat | T2T non-reference human YAO maternal | https://download.cncb.ac.cn/gwh/Animals/Homo_sapiens_v1.1_GWHDQZJ00000000/GWHDQZJ000000000.genome.fasta.gz |
| YAOpat | T2T non-reference human YAO paternal | https://download.cncb.ac.cn/gwh/Animals/Homo_sapiens_ChTY001.v1.1_pat_GWHDOOG00000000/GWHDOOG000000000.genome.fasta.gz |
| Han1 | T2T non-reference human Han1 | https://bit.ly/3JylfV3 |
| CN1mat | T2T non-reference human CN1 maternal | https://genome.zju.edu.cn/Downloads/#:~:text=v0.8/CN1_mat.v0.8-...-fasta.gz |
| CN1pat | T2T non-reference human CN1 paternal | https://genome.zju.edu.cn/Downloads/#:~:text=v0.8/CN1_pat.v0.8-,-fasta,-.gz |
| chimpT2Tpri | T2T chimpanzee primary | https://genomeark.s3.amazonaws.com/species/Pan_troglodytes/mPanTro3/assembly_curated/mPanTro3.pri.cur.20231122.fasta.gz |
| bonoboT2Tpri | T2T bonobo primary | https://genomeark.s3.amazonaws.com/species/Pan_paniscus/mPanPan1/assembly_curated/mPanPan1.pri.cur.20231122.fasta.gz |
| gorillaT2Tpri | T2T gorilla primary | https://genomeark.s3.amazonaws.com/species/Gorilla_gorilla/mGorGor1/assembly_curated/mGorGor1.pri.cur.20231122.fasta.gz |
| orangutanT2Tpri | T2T Sumatran orangutan primary | https://genomeark.s3.amazonaws.com/species/Pongo_abelii/mPonAbe1/assembly_curated/mPonAbe1.pri.cur.20231205.fasta.gz |
| borangutanT2Tpri | T2T Bornean orangutan primary | https://genomeark.s3.amazonaws.com/species/Pongo_pygmaeus/mPonPyg2/assembly_curated/mPonPyg2.pri.cur.20231122.fasta.gz |
| gibbonT2Tpri | T2T siamang gibbon primary | https://genomeark.s3.amazonaws.com/species/Symphalangus_syndactylus/mSymSyn1/assembly_curated/mSymSyn1.pri.cur.20231205.fasta.gz |
| chimpT2Talt | T2T chimpanzee alternate | https://genomeark.s3.amazonaws.com/species/Pan_troglodytes/mPanTro3/assembly_curated/mPanTro3.alt.cur.20231122.fasta.gz |
| bonoboT2Talt | T2T bonobo alternate | https://genomeark.s3.amazonaws.com/species/Pan_paniscus/mPanPan1/assembly_curated/mPanPan1.alt.cur.20231122.fasta.gz |
| gorillaT2Talt | T2T gorilla alternate | https://genomeark.s3.amazonaws.com/species/Gorilla_gorilla/mGorGor1/assembly_curated/mGorGor1.alt.cur.20231122.fasta.gz |
| orangutanT2Talt | T2T Sumatran orangutan alternate | https://genomeark.s3.amazonaws.com/species/Pongo_abelii/mPonAbe1/assembly_curated/mPonAbe1.alt.cur.20231205.fasta.gz |
| borangutanT2Talt | T2T Bornean orangutan alternate | https://genomeark.s3.amazonaws.com/species/Pongo_pygmaeus/mPonPyg2/assembly_curated/mPonPyg2.alt.cur.20231122.fasta.gz |
| gibbonT2Talt | T2T siamang gibbon alternate | https://genomeark.s3.amazonaws.com/species/Symphalangus_syndactylus/mSymSyn1/assembly_curated/mSymSyn1.alt.cur.20231205.fasta.gz |

We generated 4 genome-wide alignments referenced on hs1, chimpT2Tpri, bonoboT2Tpri, and gorillaT2Tpri, respectively. To reduce runtimes, each reference genome was first window-masked. Starting from unmasked assemblies, all windows with a score above 110 were masked (-t_thres 110)^87^. We generated local pairwise alignments with LASTZ^41^. We used the human-chimp.v2 scoring matrix with parameters (O=600 E=150 T=2 M=254 K=4500 L=4500 Y=15000)^88^. We then chained the local alignments together using kentUtils: axtChain^89^. We took several additional steps to prevent and remove misalignments during chaining. We chained alignments in each gapless region of the genome independently and only considered gapless regions greater than 1 Mb of the human genome and greater than 20 kb for each of the query genomes. This filtering allowed us to ensure a large genomic context to better separate orthologs from paralogs. We also generated a custom scoring matrix and gap penalty function^27^ for the axtChain program to more conservatively chain local alignments by preventing the chaining of alignments spanning large gaps in the target or query. We retained chains with a minimum score of 60,000 (approximately 20 kbp of matches), used kentUtils:chainNet to generate preliminary alignment nets, then refiltered chains (again at a threshold of 60,000) in the resulting net, and then ran kentUtils: chainNet to generate the final pairwise alignments for each alignable position of the reference genome. We generated multi-species genome-wide alignments with MultiZ^90^ and converted the output into an aligned FASTA file (gonomics: mafToFa).

#### Fitting a neutral model

For each of the 4 lineages analyzed for highly divergent regions, we inferred the sequence of the ancestral node in a conservative fashion. Specifically, we used the gene annotations made by NCBI RefSeq for hs1 (https://hgdownload.soe.ucsc.edu/goldenPath/hs1/bigZips/genes/hs1.ncbiRefSeq.gtf.gz) to extract fourfold degenerate codon sites and estimate branch lengths for a fixed-topology tree using a Jukes-Cantor model of evolution by maximum likelihood (PHAST: phyloFit)^91,92^. Below, we present the resulting phylogeny: ((((((humanT2T:0.000435156,(HG002mat:0.00102161,HG002pat:0.000477313)HG002anc:0.000673322)EURa nc:0.000910852,(Han1:0.000470491,((CN1mat:0.000681227,CN1pat:0.000424938)CN1anc:0.000913969,(YA Omat:0.000565216,YAOpat:0.00100586)YAOanc:0.000811958)CHNdipanc:0.000659018)CHNanc:0.00057309 3)HUMANanc:0.00606109,((chimpT2Tpri:0.000514677,chimpT2Talt:0.000464187)chimpT2Tanc:0.00153813,(bonoboT2Tpri:0.000575461,bonoboT2Talt:0.000886081)bonoboT2Tanc:0.00163245)cbaT2T:0.00445874)hcaT 2T:0.00227545,(gorillaT2Tpri:0.000928497,gorillaT2Talt:0.00102789)gorillaT2Tanc:0.00856474)hgaT2T:0.0110 735,((orangutanT2Tpri:0.00115159,orangutanT2Talt:0.00123407)orangutanT2Tanc:0.000658367,(borangutanT 2Tpri:0.000978752,borangutanT2Talt:0.00100708)borangutanT2Tanc:0.000857815)obaT2T:0.0176608)hoaT2 T:0.0116854,(gibbonT2Tpri:0.000825439,gibbonT2Talt:0.00102365)gibbonT2Tanc:0.0116854)hgiaT2T;

#### Ancestral state inference

For each position in the alignment, a base was determined to be present in the ancestral node according to the tree, i.e., a base is present in at least two species on two independent lineages connected to the ancestral node (lineages to parent node and two child nodes). This is equivalent to treating aligned bases as having a common origin. For bases that are present at a node, we reconstructed the probabilities of A, C, G, and T in the ancestral node using the estimated branch lengths and the value of the base position in extant species ^93^. Rather than assigning a single deterministic base to the ancestral node from these 4 probabilities, we stored the vector of probabilities of A, C, G, and T for each base (gonomics:reconstructSeq).

#### Quantifying divergence

To identify regions of significant divergence between the inferred ancestral nodes (e.g. human-chimpanzee common ancestor and human ancestral node), we counted the number of mutations or divergences separating the nodes. We counted each gap separating nodes as one divergent event, irrespective of length.

To measure substitutions between ancestral nodes, we first calculated the cosine distance between the inferred 4-state probability vectors corresponding to the posterior probability of each possible base at that ancestral node. This metric represents the probability that a substitution occurred on the branch separating these two ancestral nodes. We then binarized sites to either substitutions (cosine distance ≥ 0.8) or non-substitutions (cosine distance < 0.8), to conservatively estimate divergence. We then quantified the divergence of a genomic region as the sum of the number of substitutions and gaps in that region.

We implemented a program to count the number of divergences that separate the inferred ancestral node and the consensus node of the extant species by sliding along the genome alignment in windows of 500 bp (gonomics: pfaFindFast). Windows with significantly more divergence than expected were identified as HAQERs, chimp-AQERs, bonobo-AQERs, and gorilla-AQERs. To conservatively estimate significance, rather than use the genome-wide average divergence rate, we used the maximum divergence rate in a 10 Mbp window of the genome as the estimate of the expected divergence rate when defining HAQERs. For the other lineages, we scaled this conservative divergence rate relative to the ratio of branch lengths between the human branch and the branch length of the other lineages from fourfold degenerate sites for: chimp-AQERs, bonobo-AQERs, and gorilla-AQERs. The p-value of obtaining at least the number of divergences found in each 500 bp window along the assembly was calculated using a binomial distribution, where the number of trials was 500, and the probability of success was the expected divergence rate. This raw p-value was adjusted using the Benjamini-Hochberg procedure by counting the number of 500 bp windows in the genome and ranking their raw p-values. Windows with at least 29 mutations in a 500 bp were identified as HAQERs, the same threshold we previously used^27^. This was equivalent to an adjusted p-value of less than 3e-7 for T2T and Consensus HAQERs (1.5e-7 for Gapped HAQERs). Applying this adjusted p-value threshold to all lineages, the mutation threshold in a 500 bp window was 29 for chimp-AQERs and bonobo-AQERs, and 36 for gorilla-AQERs, due to the longer phylogenetic branch from the human-gorilla ancestor to gorilla (gonomics: bedFormat, gonomics: bedFilter, gonomics: mergesort, gonomics: bedMerge).

### AQER subsets

As previously described, we generated several subsets of 500 bp window size Consensus HAQERs^32^.

We generated a repeat-free HAQER subset excluding centromeres/satellites, segmental duplications, simple repeats, or RepeatMasker annotations on hs1^32^. We defined a repeat subset of HAQERs as the complement of the repeat-free subset. We also generated high-divergence HAQER subsets with mutation thresholds at 34 and 39 mutations, elevated from the original threshold of 29 mutations.

### Genomic regions of interest

Genomic regions of evolutionary interest analyzed in this study are presented in the table below.

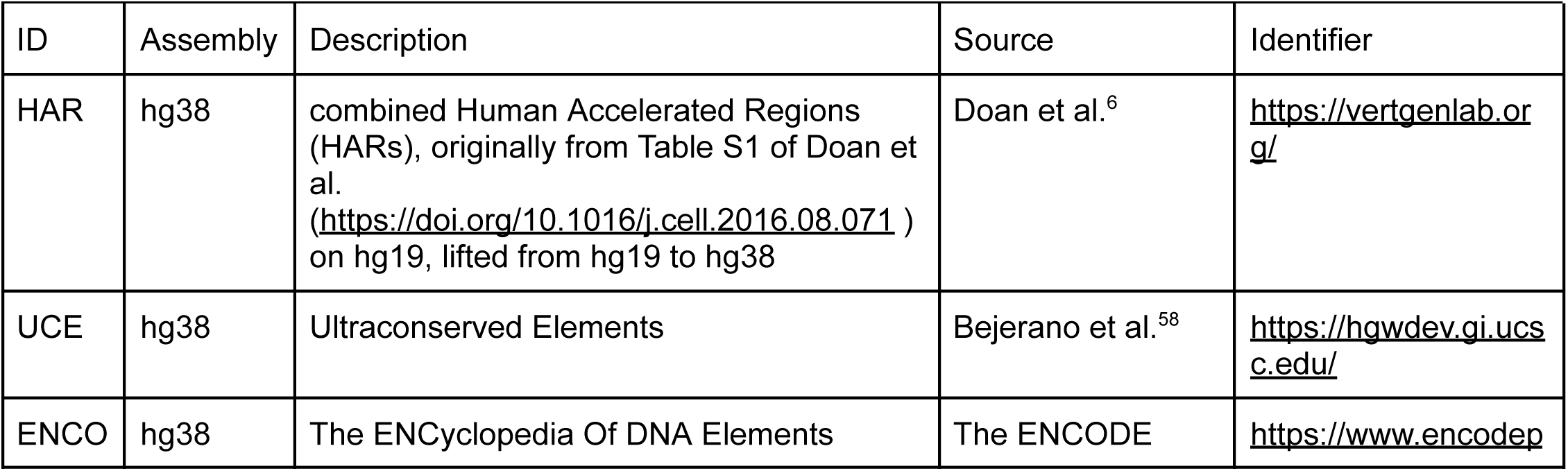
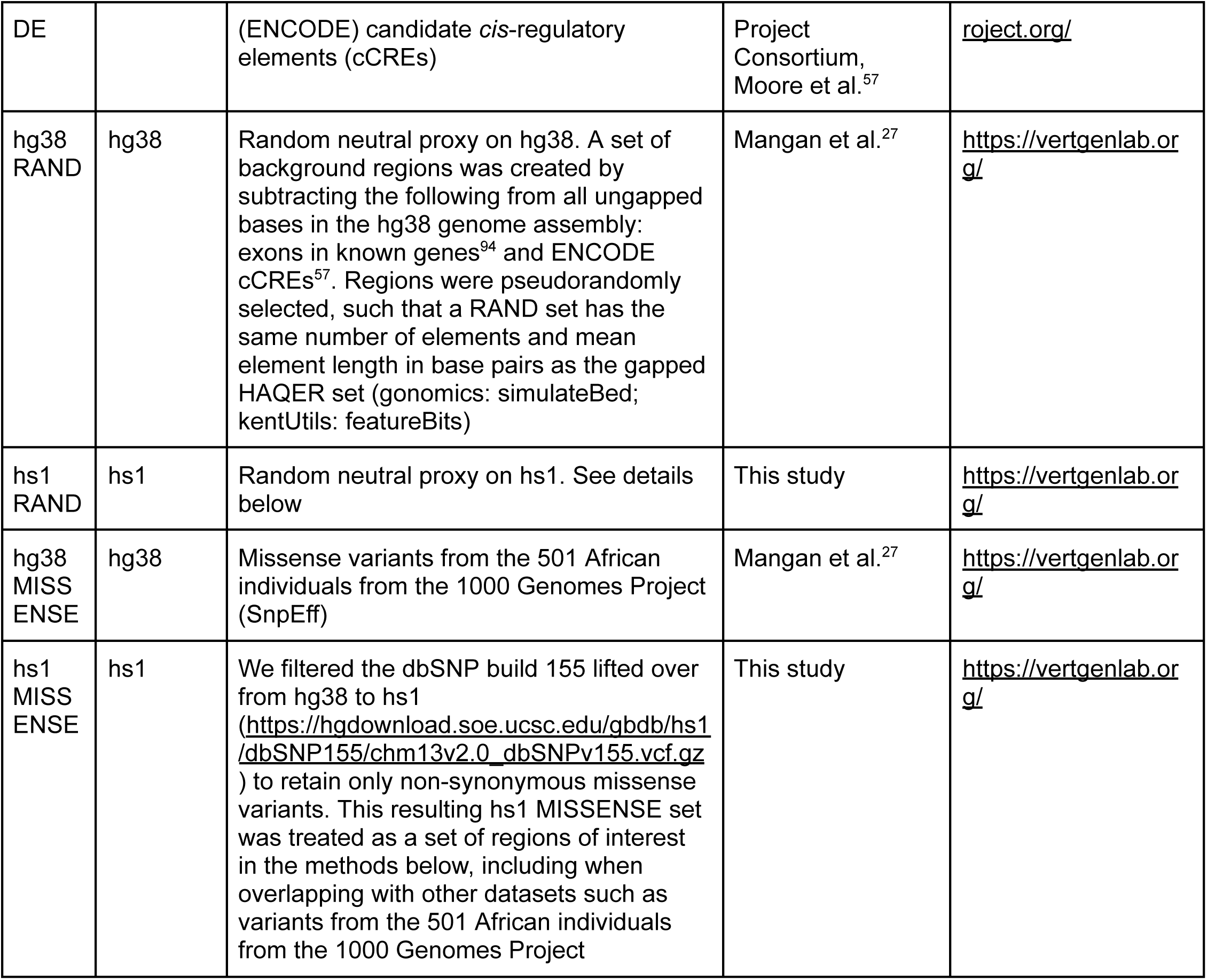

To generate hs1 RAND, or neutral proxy random genomic regions on the hs1 genome, we pseudorandomly selected elements from a set of background regions excluding functional annotations. Specifically, we used bedtools:subtract^95^ to exclude ENCODE cCREs (lifted from hg38 to hs1; see ‘Lifting regions across genome assemblies’), exons in known genes (accessed from the following URL: https://s3-us-west-2.amazonaws.com/human-pangenomics/T2T/CHM13/assemblies/annotation/chm13v2.0_RefSeq_Liftoff_v5.2.gff3.gz) with 50 bp bidirectional padding (bedtools:slop), conserved regions, and regions inaccessible to alignment. Conserved regions were compiled from the following sources: hg38 30-primate PHAST by the UCSC Browser Group (https://hgdownload.soe.ucsc.edu/goldenPath/hg38/database/phastConsElements30way.txt.gz) lifted to hs1 using methods described below (see ‘Lifting regions across genome assemblies), hs1 PHAST(https://github.com/marbl/T2T-Browser?tab=readme-ov-file)^32,96^, hs1 conserved regions identified by following the AcceleratedRegionsNF pipeline and implementing 2 sets of PHAST parameters, “target_coverage: “0.40” and “expected_length: “45””” and “target_coverage: “0.35” and “expected_length: “40””” (https://github.com/keoughkath/AcceleratedRegionsNF). Regions inaccessible to alignment were defined as regions with insufficient information to infer the T2T human-chimpanzee ancestor sequence at ≥ 80% confidence (gonomics: reconstructSeq, gonomics: multiFaScan, gonomics: bedMerge). This background occupies approximately 80% of the bases of the hs1 genome assembly. From the background, regions were pseudorandomly selected, such that a RAND set has the same number of elements and mean element length in base pairs as the T2T HAQER set (gonomics: simulateBed). Three replicate hs1 RAND sets were generated from 3 pseudorandom seeds (10, 40, 70). The set with pseudorandom seed 10 served as the reference set in statistical tests.

### Lifting regions across genome assemblies

Regionss were lifted from hg38 to hs1 and vice versa on the UCSC Genome Browser LiftOver web interface (https://genome.ucsc.edu/cgi-bin/hgLiftOver) using the parameter-minMatch=0.7 (minimum ratio of bases that must remap), not allowing multiple output regions. We then merged overlapping entries (gonomics: bedMerge).

#### GC content analysis

We calculated the GC-content in each 500 bp window of hs1 along the genome alignment (gonomics: faFindFast). We also implemented a program to calculate the GC-content of each HAQER (gonomics: gcContent). To create GC-content density and bar plots, we randomly sampled 1% of the 500 bp windows from the whole genome (gonomics: bedFilter). We plotted the percentage of GC-rich (GC > 0.75) and GC-poor (GC < 0.25) regions and assigned significance via the Chi-squared test of independence against the observed percentages of GC-rich and GC-poor regions in the whole genome. For each region set, X, a 2×2 contingency table was constructed with the dimensions {X, Whole Genome} and {GC-rich, Non-GC-rich} or {GC-poor, Non-GC-poor}. We conducted this analysis on hs1. Gapped HAQERs were lifted from hg38 to hs1 (see Lifting regions across genome assemblies).

### Incomplete lineage sorting

To analyze the relationship between Consensus HAQERs and regions of incomplete lineage sorting (ILS), we accessed genome-wide posterior probability distributions of ILS states inferred by the TRAILS (tree reconstruction of ancestry using incomplete lineage sorting) hidden Markov model^43^. TRAILS infers four hidden states (V0-V3) representing canonical and non-canonical great ape topologies. We calculated two-sided binomial-based overlap enrichments between Consensus HAQERs and regions of the human genome with a posterior probability of at least 0.6 for each TRAILS hidden state (gonomics:overlapEnrichments). Significance was assigned for enrichment and depletion at Bonferroni-adjusted p < 0.05.

### Repeat enrichment analysis

We obtained RepeatMasker and segmental duplication annotations developed natively for each T2T genome assembly^32^. RepeatMasker annotations were split into multiple files by repeat class (gonomics: bedSplit, gonomics: bedMerge). We calculated the overlap enrichment and depletion between each repeat class and each AQER set (gonomics: overlapEnrichments). We reported enrichment as the ratio between the observed number of overlaps and the expected value of overlaps. We calculated Bonferroni-adjusted p values for enrichment and depletion. Significance was assigned for enrichment and depletion at pAdj < 0.05.

### Mutation rate estimation and comparison

We obtained mutation rate estimates for autosomes from Seplyarskiy et al.^56^. For each position in the genome with mutation rate (MR) estimates, we summed the MRs of the three possible substitutions (e.g., for an A reference base, the rates of A→C, A→T, and A→G) to calculate the total MR, representing the probability of mutating to any other base. We overlapped the processed MR estimates with each region set (gonomics: intervalOverlap). For each element, we calculated the mean of the total MRs across all overlapping positions. The mean MRs of each region set were then compared against those of the hg38 RAND with Bonferroni-adjusted pairwise t tests.

We conducted this analysis on the hg38 assembly. To enable comparison, the following region sets were lifted from hs1 to hg38 (see ‘Lifting regions across genome assemblies): T2T HAQERs, Human Common Ancestor HAQERs, hs1 RAND.

### Human genetic variation preprocessing

We obtained hs1-aligned haplotype-phased genotype data from 2,504 human samples gathered by the 1000 Genomes Project^97,98^, which are available at the following url: https://github.com/JosephLalli/phasing_T2T?tab=readme-ov-file. We filtered the genotype data to retain only biallelic substitution variants that segregate in the 501 African individuals, reducing the impact of population bottlenecks introduced by Out-of-Africa migration events (gonomics: vcfFilter). We also created filtered genotype files for each of the 5 African populations (Gambian in Western Division – Mandinka, Mende in Sierra Leone, Esan in Nigeria, Yoruba in Nigeria, and Luhya in Webuye, Kenya) containing only individuals from that population and variants that segregate within that population.

We retained both polarized and unpolarized versions of the filtered genotype data. To create the polarized version of the filtered genotype data, we first constructed an alternate hs1 sequence where the reference allele at each segregating site is replaced with the alternate allele (gonomics: vcfToFa) and appended the alternate hs1 sequence to our multiple alignment. Then, we reconducted ancestral state inference, using the expanded alignment and treating hs1 and alternate hs1 as equal-weight branches of an “hs1 consensus haplotype” (gonomics: reconstructSeq). Unlike when we identified the T2T HAQER set^32^, we did not bias the ancestral state inference toward hs1. Instead, from the 4 base probabilities for the human-chimpanzee ancestor, we accepted the most likely allele as the ancestral state if its probability was greater than or equal to 99%. Otherwise, we assigned an N to the ancestral state, to ensure that only high confidence SNPs were retained for the subsequent analysis of derived allele frequencies. Using the newly inferred human-chimpanzee ancestor sequence, we annotated the ancestral allele for each variant (gonomics: vcfAncestorAnnotation) and removed polymorphic sites where neither the reference allele nor the alternate allele present in the extant human population matched the allele present in the inferred T2T human-chimpanzee ancestor sequence.

### Allele frequency spectra

We generated allele frequency spectra on hs1 and hg38. We created subsets of filtered African variants (see ‘Human genetic variation preprocessing’) that overlap ROIs (gonomics: intervalOverlap). To limit the impact of linkage disequilibrium on the shape of the derived allele frequency spectrum, we retained variants that were a minimum of 10,000 bases away from any other variant in the sample set (gonomics:proximityBlockVcf). We used 3 pseudorandom seeds for sampling. We then calculated allele frequency spectra (gonomics: vcfAfs). We generated allele frequency spectra with both polarized and unpolarized variants. In the polarized version, we measured derived allele frequencies (DAF). In the unpolarized version, we measured minor allele frequencies (MAF), which ranged between 0 and 0.5.

For this analysis, we filtered the ENCODE cCRE set to retain only those with Z-score > 5, which corresponds to above 99th percentile for a one-sided Gaussian distribution DNase-seq and at least one of 3 ChIP-seq marks (H3K4me3, H3K27ac, and CTCF).

For the analysis on hs1, the following region sets were lifted from hg38 to hs1 (see ‘Lifting regions across genome assemblies’): Gapped HAQERs, HARs, ENCODE, UCEs, hg38 RAND.

For the analysis on hg38, the following region sets were lifted from hs1 to hg38 (see ‘Lifting regions across genome assemblies’): T2T HAQERs, Consensus HAQERs, hs1 RAND.

### Selection parameter estimate

To infer the direction and magnitude of selective pressure acting on region sets, we used the previously described hierarchical Bayesian model implementing the Metropolis-Hastings algorithm for Markov Chain Monte Carlo (MCMC) sampling^27^ to estimate mean selection parameters of each region set from their polarized proximity blocked human population variant allele frequency data by subpopulation (see ‘Allele frequency spectra’), applying divergence-based ascertainment corrections^99^ to the following region sets: HAQERs, HARs, UCEs (gonomics: selectionMcmc). We then calculated the mean and 95% highest density credible interval for each Marokv chain, discarding the first 5,000 iterations as burn-in for variant sets overlapping regions of evolutionary interest (gonomics: mcmcTraceStats). The mean selection parameter μ of a region set should be μ ≈ 0 for regions under neutral selection, with μ < 0 and μ > 0 indicating negative and positive selection, respectively.

### Simulated genetic variation

We generated simulated genetic variation datasets corresponding to Consensus HAQERs and HARs, each containing 1,002 alleles. For each region set, the number of segregating sites and the selection parameter were set to the mean values observed after overlapping filtered polarized African variants and applying proximity blocking. Simulated variant calls were generated using gonomics:simulateVcf, and synthetic allele frequency spectra were calculated with gonomics:vcfAfs. All simulations were performed on the hs1 genome assembly.

### Fixed divergent allele proportion analysis

We define “divergent” sites as positions where the human ancestral node and human-chimpanzee ancestor node disagree (see ‘Quantifying divergence’) and “fixed” sites as divergent positions with no observed polymorphism in modern human populations. To prevent uneven gap sizes from skewing the overall trend, we represent divergence with substitutions only, excluding gaps. To calculate the proportion of fixed divergent alleles within each region set, we first identified all divergent positions between hs1 and the inferred T2T human-chimpanzee ancestor (gonomics:multiFaToVcf). We then identified divergent positions overlapping each region set (gonomics: intervalOverlap). We intersected the set of divergent positions with the set of all polymorphic sites identified in the 501 African individual subset of the hs1 filtered 1000 Genomes Project data (see ‘Human genetic variation preprocessing’) (gonomics: intervalOverlap). Positions found in both sets were labeled as polymorphic divergent sites, and divergent sites not found in the 1000 Genomes Data were labeled as fixed divergent sites. We then determined the sets of fixed and polymorphic divergent sites overlapping each region set (gonomics: intervalOverlap). We plotted the proportion of divergent sites that are polymorphic as the number of polymorphic sites divided by the sum of polymorphic and fixed divergent sites and assigned significance via the Chi-squared test of independence against the observed ratio of polymorphic divergent sites in hs1 RAND. The 2×2 contingency table for this analysis for a set of regions, X, had the dimensions {X, hs1 RAND} and {Fixed, Polymorphic}. For the analysis on hs1, the following region sets were lifted from hg38 to hs1 (see ‘Lifting regions across genome assemblies’): gapped HAQERs, HARs, ENCODE, UCEs, hg38 RAND. We also conducted the analysis on hg38 using previously described methods, lifting over the following region sets from hs1 to hg38 (see ‘Lifting regions across genome assemblies’): T2T HAQERs, Human Common Ancestor HAQERs, hs1 RAND.

### Polymorphism and divergence comparative analysis

Using the set of all divergent sites that overlap each region set (see ‘Fixed divergent allele proportion analysis’), we calculated the divergent sites per base as the number of divergent sites divided by the total length in base pairs of the input region set. Similarly, we calculated polymorphic sites per base as the number of variants from the 501 African individual subset of the hs1 filtered 1000 Genomes Project data (see ‘Human genetic variation preprocessing’) that overlapped each region set divided by the length in base pairs of that set. We plotted the polymorphic sites per base against the divergent sites per base for each region set.

### Polymorphism and divergence distribution analysis

To analyze the distribution of polymorphism and divergence in the human genome, we computed counts of divergent sites and polymorphisms in sliding windows across autosomes using our genome alignment. We used two window sizes: 50 kb and 500 bp. We chose 50 kb instead of a smaller window size to minimize the contribution of noise^100^. To prevent uneven gap sizes from skewing the overall trend, we represent divergence with substitutions only, excluding gaps. Divergent substitution counts were obtained by comparing the inferred human-chimpanzee ancestor to the human ancestral node (gonomics:pFaFindFastExpanded). Polymorphism counts were obtained by overlapping windows with polymorphic sites identified in the 501 African individual subset of the hs1 filtered 1000 Genomes Project Data (see ‘Human genetic variation preprocessing’) using bedtools:intersect^95^.

For each window, we recorded the number of high-confidence human-chimpanzee ancestor bases, where a base was considered confident if its posterior probability was at least 0.8. For downstream visualization, only windows with confidence per base exceeding 0.9 were retained. From the 50 kb windows, we randomly sampled 1% genome-wide. From the 500 bp windows, we retained all windows fully contained within HAQERs (GenomicRanges^101^) and sampled 1% of the remaining windows. We plotted polymorphisms per base against substitutions per base and constructed heatmaps with a bin width of 0.001 x 0.001 for the 50 kb window plot and 0.002 x 0.002 for the 500 bp window plot. We excluded bins with no more than 50 counts, which may be artifact-heavy.

### Chromatin state analysis

We analyzed chromatin state enrichments using ChromHMM^102^ annotations from two resources: the 15-state model across 127 epigenomes from the Roadmap Epigenomics Consortium^62^ and the 18-state model across 833 epigenomes from EpiMap^17^. We calculated overlap enrichments and depletions between each HAQER set and each chromatin state using a two-sided binomial-based statistical framework (gonomics:overlapEnrichments). Enrichments are reported as the ratio of observed to expected overlaps, and Bonferroni-adjusted p values were used to assess significance, with pAdj < 0.05 considered significant for both enrichment and depletion.

To investigate which HAQERs overlap bivalent regulatory states, we identified the number and identity of EpiMap epigenomes annotated with bivalent enhancers or bivalent promoters that overlapped each Consensus HAQER using bedtools:intersect^95^.

These analyses were conducted on hg38. The following HAQER sets and subsets were lifted from hs1 to hg38 (see ‘Lifting regions across genome assemblies’): T2T HAQERs, Consensus HAQERs, repeat-free, repeat, GC content > 0.75, GC content < 0.25, ≥31 mutations, ≥34 mutations, ≥39 mutations, window size HAQERs.

### GWAS catalog trait enrichment analysis

We retrieved GWAS variants from the GWAS catalog^103^ (gwas_catalog_v1.0.2-associations_e113_r2025-01-30.tsv) and filtered to retain only those segregating in the GBR subpopulation of the 1000 Genomes Project^104^. To expand to linked variants^105^, we used plink –-r2 to identify all 1000 Genomes Project variants in linkage disequilibrium with each GWAS variant (Plink R^2^ > 0.7). For each trait mapped to an Experimental Factor Ontology, we merged the set of GWAS variants with their linked variants and tested for enrichment in HAQERs using gonomics:overlapEnrichments, assigning significance at FDR-adjusted p < 0.05.

This analysis was conducted on hg38. Consensus HAQER and T2T HAQER sets were lifted from hs1 to hg38 (see ‘Lifting regions across genome assemblies’). On hg38, gapped HAQERs that do not have any bases overlapping T2T HAQERs and T2T HAQERs that do not have any bases overlapping gapped HAQERs were combined to form a set of variable HAQERs (gonomics: intervalOverlap –nonOverlap).

### scSTARR-seq (single-cell Self-transcribing active regulatory region sequencing)

#### Input library preparation

We used scSTARR-seq (single-cell Self-transcribing active regulatory region sequencing) to assess the regulatory activity of HAQERs. We synthesized test sequences and cloned them into the STARR-seq screening vector using a commercial service (Twist Bioscience). Plasmids were transformed into One Shot Stbl3 chemically competent E.coli (Thermo Fisher), selected for ampicillin resistance, and amplified in Luria broth (Invitrogen) with 100 μg/mL ampicillin. Endotoxin-free plasmids were then purified with the ZymoPURE II Plasmid Maxiprep kit per manufacturer’s instructions (Zymo Research). As necessary, purified plasmids were then precipitated with 3M Na-Acetate pH 5.2, and 100% ethanol for 2 hours to achieve desirable concentrations.

We prepared a solution containing each STARR-seq plasmid at a total plasmid concentration of 3 μg/mL for *in vivo* scSTARR-seq and 1.6 μg/mL for *in vitro* scSTARR-seq. The pooled STARR-seq solution was then mixed with a pCAG-GFP injection reporter plasmid. The pCAG-GFP injection reporter plasmid represented ⅙ and ⅓ of the total plasmid content of the final input library solution in *in vivo* scSTARR-seq and *in vitro* scSTARR-seq, respectively.

Our *in vivo* STARR-seq input library included plasmids with a total of 75 distinct inserts: 8 HAQER sequences, 63 sequences of interest for other projects, and 4 scrambled pseudorandom sequences which served as negative controls. The 4 pseudorandom sequences with the lowest enhancer activity in past trials were chosen^27^.

Our *in vitro* STARR-seq input library included plasmids with a total of 77 distinct inserts: 8 HAQER sequences, 65 sequences of interest for other projects, and the same 4 negative control sequences in the *in vivo* library.

We performed input normalization as previously described^27^.

#### *In utero* electroporation

For *in vivo* scSTARR-seq, we delivered input libraries via *in utero* electroporation. E14.5 wild type CD1 pregnant females were anesthetized with isoflurane. Uterine horns were exposed by abdominal incision. Each embryo was injected with 1-1.5 μl of plasmid solution (containing 0.01% fast green and 1-2 μg/μl of plasmids) and electroporated using platinum-plated BTX Tweezertrodes with five 50 ms-pulses at 50V with 950 ms pulse-interval. Uterine horns were then repositioned into the abdominal cavity and the muscle and skin incisions were sutured. Dams were then placed on a heating pad for recovery and monitored.

We harvested electroporated brains after approximately 48 hours. Then, the harvested brains were dissected in ice-cold sterile PBS. Meninges were removed and GFP+ portions of the cortices were incubated at 37◦C for 10 minutes in 0.25% trypsin-EDTA supplemented with 0.1% DNAse I (New England Biolabs cat# M0303S).

Following incubation, the trypsin solution was removed and replaced with ice-cold 10% FBS/HBSS/Propidium iodide (Invitrogen) supplemented with 0.01% DNAse I. A single cell suspension was then generated by trituration with a fire-polished glass pipette and filtered with a 30 μm cell strainer.

#### *In vitro* transfection

For *in vitro* scSTARR-seq, we differentiated hiPSC (human induced pluripotent stem cells) of the DU11 cell line into cardiomyocytes as previously described^106,107^. We delivered the input library via transfection after at least 30 days of differentiation, following manufacturer protocols for Viafect (Promega). We incubated the input library for 2 days. At the end of the incubation period, we detached the cardiomyocytes with Accutase to create a single cell suspension.

#### FACS (Fluorescence activated cell sorting)

We stained harvested cells with the LIVE/DEAD Near-IR Dead Cell Stain per manufacturer’s instructions (Thermo Fisher). Following staining, viable GFP+ cells were bulk sorted using a FACS Aria II cytometer (BD Biosciences).

#### scSTARR-seq output library preparation, sequencing, and preprocessing

Up to 10,000 GFP+ cells were captured per lane of a 10X Chromium device and single cell output libraries were prepared using protocols from the Chromium Next GEM Single Cell 3′ Reagent Kits v3.1 (Rev D) User Guide (10X Genomics, Inc.). Final libraries were quantified using the Bioanalyzer (Agilent) according to the manufacturer’s protocols. Prior to enzymatic fragmentation, an aliquot of cDNA was separated for targeted enrichment. Targeted enrichment was conducted identically to previously reported^27^.

Final 10X libraries were sequenced using NextSeq 2000 reagents (Illumina; R1 28, I1 10, I2 10, R2 90). We sequenced reporter-targeted enrichment libraries independently on the Illumina NextSeq 1000 platform. FASTQ files from both output libraries were recovered with BCL Convert (v3.10.4, Illumina). We used the following bases mask: Y28;I10;I10;Y90. Output FASTQ files from both the unenriched and reporter-targeted enrichment libraries across multiple lanes were concatenated together for downstream analysis.

For *in vivo* scSTARR-seq, GFP+ cells were pooled from all embryos in each experiment to control for batch effects associated with anatomical differences in electroporation and dissection. We sequenced 17,564 single cells from the output library, which was composed of GFP+ cells pooled from 5 embryos from a single mouse injected at E14.5 and harvested at E16.5.

For *in vitro* scSTARR-seq, we sequenced 5,648 single cells.

#### Enhancer activity quantification

To score enhancer activity from scSTARR-seq data, we produced count matrices using CellRanger v6.0 (10x Genomics) with a custom reference genome. For *in vivo* scSTARR-seq, the custom reference genome includes the sequence of the mouse reference mm10 with additional FASTA records containing the sequences of each STARR-seq reporter construct and the pCAG-GFP injection reporter sequence. For *in vitro* scSTARR-seq, the custom reference genome includes the sequence of the human reference hg38 with additional FASTA records containing the sequences of each STARR-seq reporter construct and the pCAG-GFP injection reporter sequence.

For *in vivo* scSTARR-seq, we performed subsequent analysis for cluster identification in Seurat v4.0^108^. Cells containing 200 or fewer genes or more than 7,500 genes were removed. The library was normalized, and 2,000 highly variable features were identified. Variation in gene expression based on cell-cycle related genes was regressed from cluster analysis in dataset scaling using an annotated set of G2M and S phase related genes provided in Seurat. k-nearest neighbors (k=20) were calculated in the space of significant principal components (in this case, 30 principal components) and clustering was performed with the Louvain-Jaccard method. Visualizations were generated in UMAP (uniform manifold approximation and projection) space^109^. Cell clusters with no GFP reads were removed, and the remaining cells were re-clustered. We manually assigned cell type identities to each cluster based on each cluster’s expression of cell type marker genes. Cell type marker genes were curated from literature^110–114^. Multiple clusters corresponding to the same cell type were pooled as metaclusters for subsequent analysis.

For both *in vivo* and *in vitro* scSTARR-seq, we implemented *gonomics: starrSeqAnalysis* to remove unique molecular identifier (UMI) duplicates and normalize the output library read count of each construct by both the input library read count of the same construct and the mean output library read count of the negative control constructs. We quantified the enhancer activity score for each construct as the input and negative control-normalized UMI count. To calculate cell type-specific enhancer activity score for each construct, we grouped input-normalized output library UMI counts by construct and cell type, then conducted negative control normalization for each cell type.

To calculate the statistical significance of the enhancer activity scores, we first determined that gamma distribution fit our negative controls better than normal distribution, based on its support for values from 0 to positive infinity, the appearance of the distribution and quantile-quantile plots, and the values of goodness-of-fit and Shapiro-Wilk normality tests. We fit a gamma distribution to the enhancer activity scores of the negative controls and calculated the right-tailed p-values of each construct. We conducted this statistical analysis both overall and by cell type.

To examine the read distribution among cells in the *in vitro* scSTARR-seq experiments, we additionally used *gonomics: starrSeqAnalysis* to create pseudoreplicates by randomly assorting cells into 1 of 5 bins. The number 5 was chosen because it was the maximum number of bins that still assured that each bin still had >1 read of each negative control construct.

### Genome Browser visualizations

We visualized several tracks on the UCSC Genome Browser on hs1, using both NCBI RefSeq and CAT/Liftoff Genes for gene annotations. To visualize divergence density between the reconstructed human-chimpanzee ancestor and the human ancestral node, we converted the BED file listing divergences for each 500 bp window into a wiggle (WIG) format track for the UCSC Genome Browser (gonomics: bedToWig). We then converted this WIG track into a binary wiggle (bigWig) format track for final visualization on the browser (kentUtils: wigToBigWig).

To visualize open chromatin data, we obtained FASTQ files from published studies^115,116^, aligned them to hs1 using the Burrows-Wheeler Aligner (BWA)^117^, and displayed the resulting BAM files as density graphs on the browser.

## Supplementary Figures

**Figure S1:**
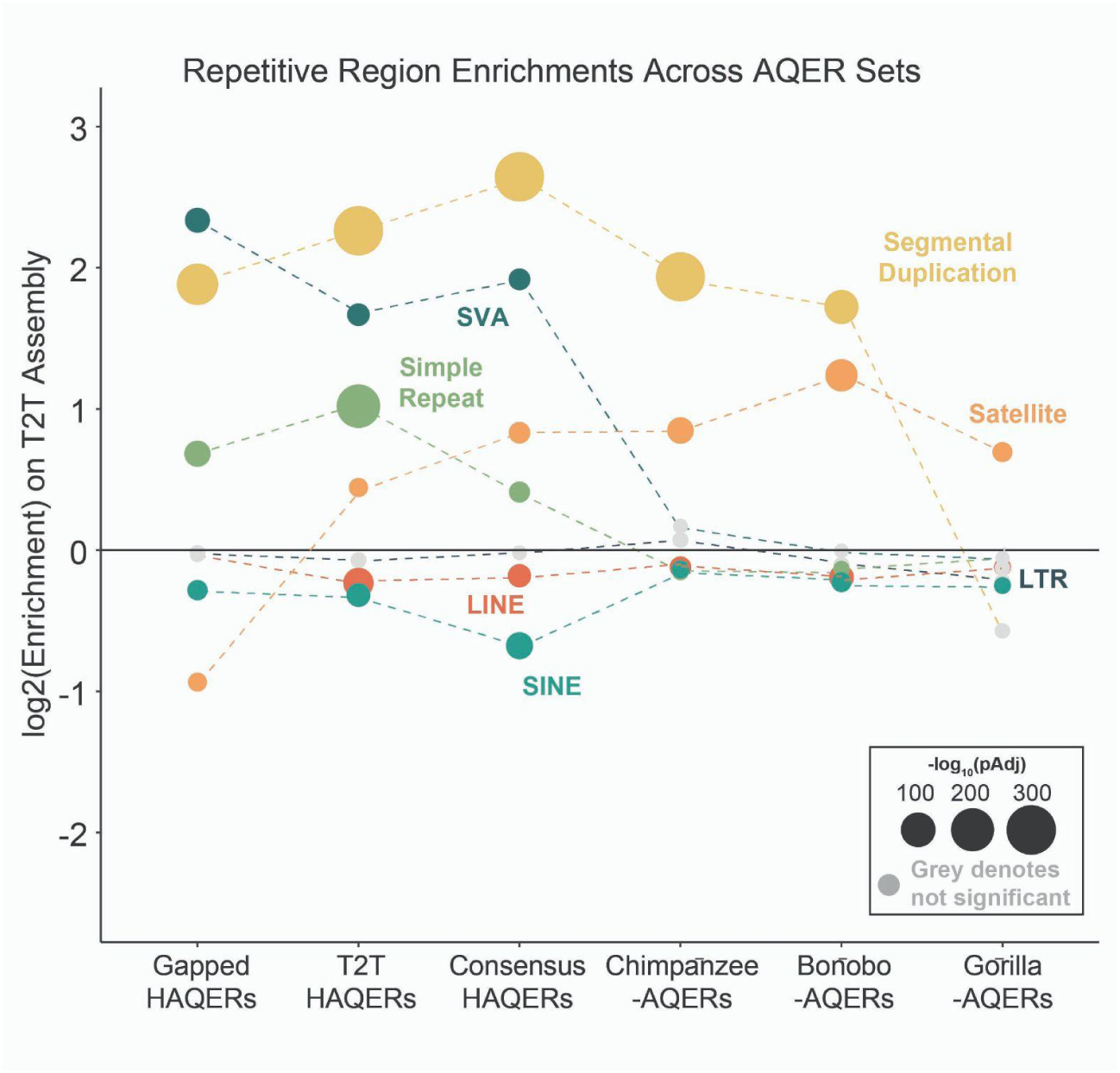
Repetitive region enrichments across HAQER and non-human primate AQER sets, related to Figure 2. Enrichment of HAQER sets and chimpanzee-, bonobo-, and gorilla-AQER sets relative to the genome in major repetitive element classes and families. Ape AQER sets show similar enrichments to HAQERs in segmental duplications and satellite DNA, suggesting similar patterns of rapid evolution across repeat classes across independent primate lineages.

**Figure S2:**
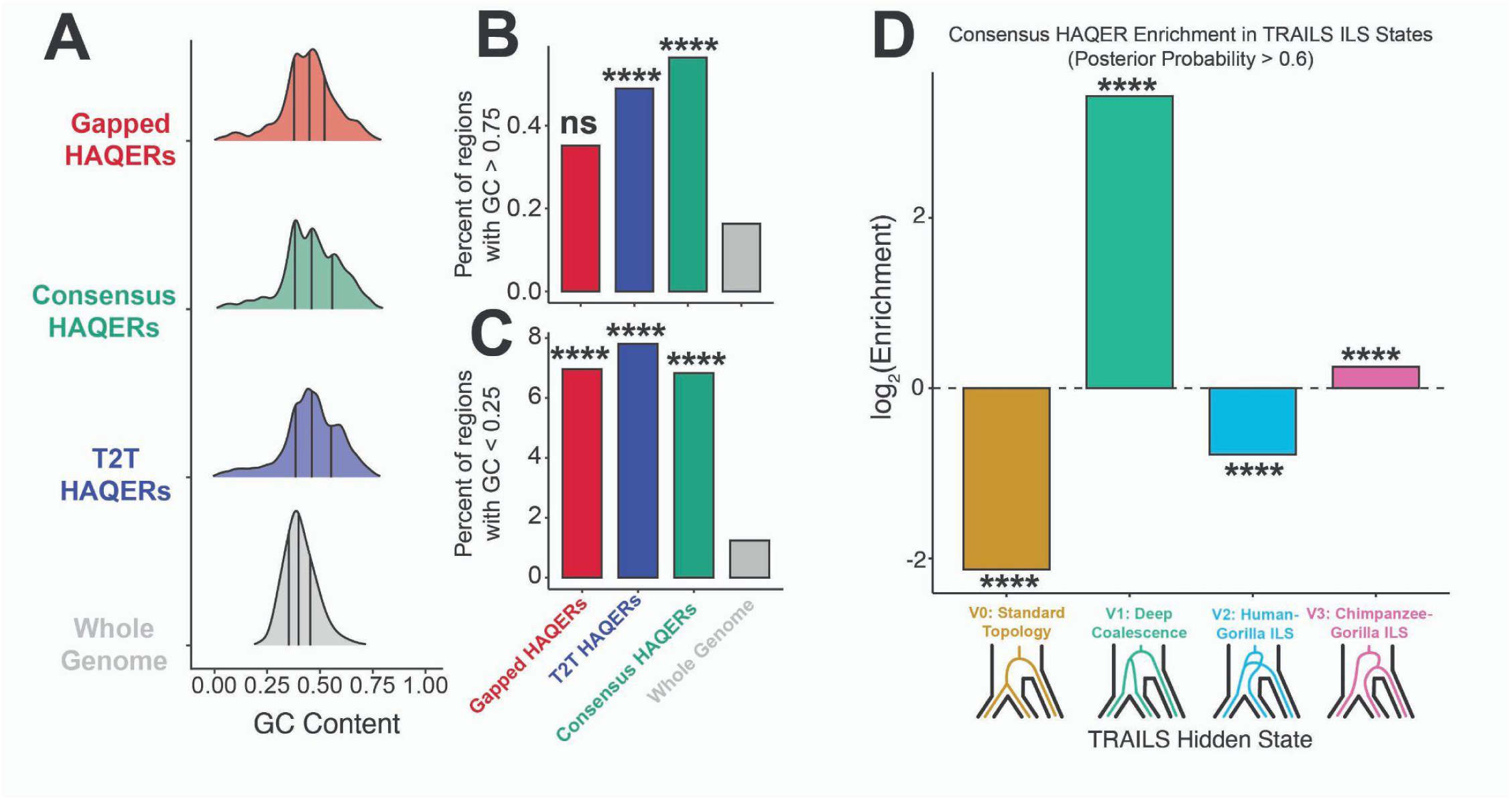
GC content and ILS states of rapidly evolved genomic regions, related to Figure 2. (A) Distribution of GC content across 3 HAQER sets (Gapped HAQERs, T2T HAQERs, Consensus HAQERs) compared with random 500 bp windows sampled from the human genome (Whole Genome). (B-C) Percentage of (B) GC-rich (GC > 0.75) and (C) GC-poor (GC < 0.25) regions across HAQER sets and the whole genome. Statistical significance was assessed by pairwise chi-squared tests against the whole genome. (D) Overlap enrichments between Consensus HAQERs and regions of incomplete lineage sorting (ILS) estimated as hidden states of the TRAILS ILS model^43^. Consensus HAQERs are significantly depleted from the V0 state (standard great ape topology) and enriched in the V1 state (standard great ape topology regions with long branches). HAQERs are depleted from the V2 nonstandard topology (human-gorilla ILS) and slightly enriched in the V3 state (chimp-gorilla ILS). (**** p < 0.0001; ns = not significant)

**Figure S3:**
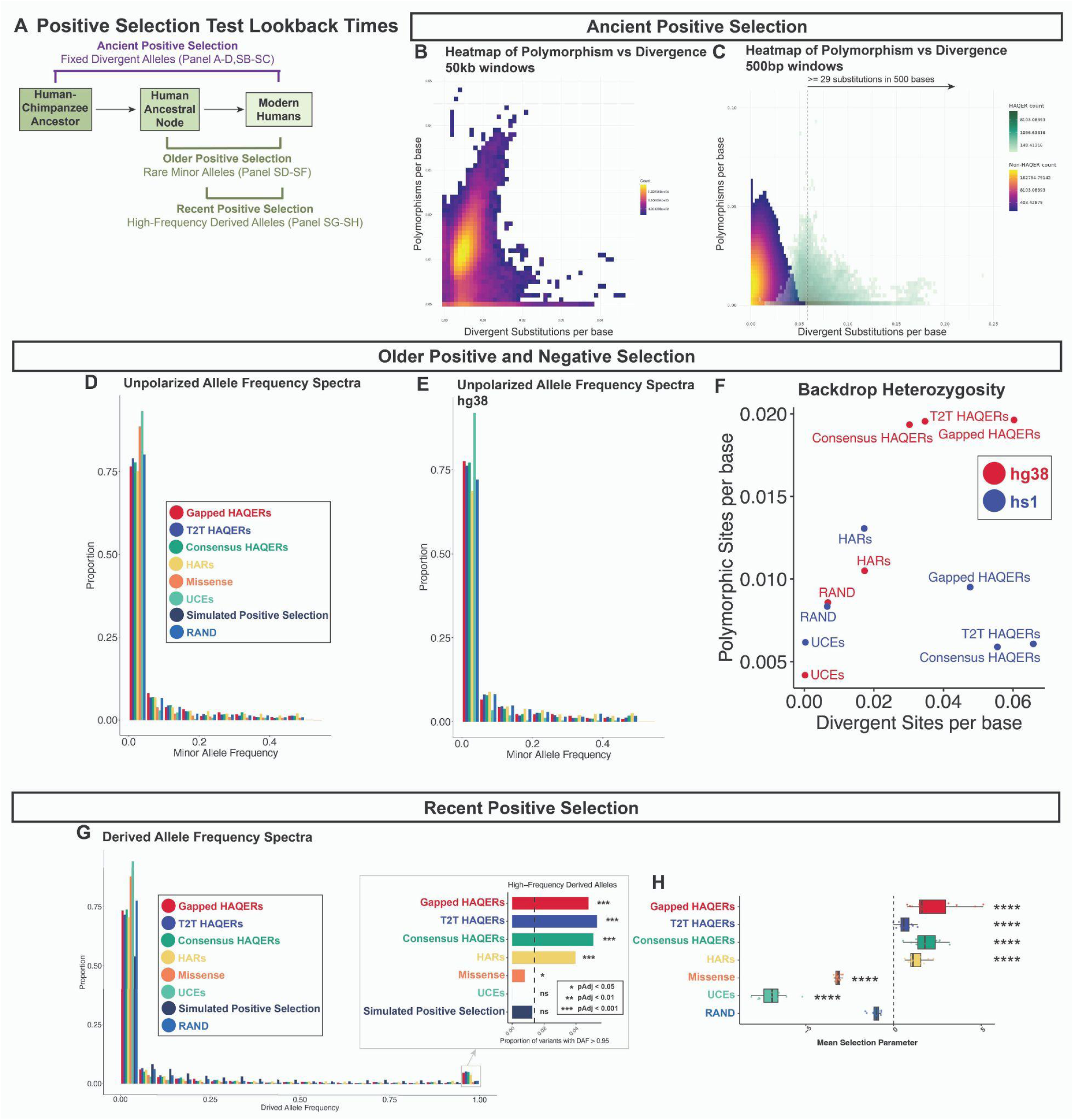
Expanded selection analysis of HAQERs, related to Figure 3. (A) Schematic comparing tests to detect positive selection at different time scales. First, the proportion of fixed divergent alleles detects “ancient” positive selection on the order of millions of years. Ancient beneficial divergences that occurred millions of years ago have enough time to complete selective sweeps, become fixed in the population, and regain polymorphism^60,61^. However, their polymorphism-to-divergence ratio would still be lower than expected due to positive selection. Second, a high proportion of rare minor alleles suggests “older” positive selection, within the past hundreds of thousands of years, enough time for derived beneficial divergences and their nearby linked variants to complete selective sweeps, become fixed in the population, and experience an initial restoration of heterozygosity. Because mutations are rare overall, they restore polymorphism in the positively selected regions slowly. Initially, mutated alleles in the positively selected regions have very low frequencies. At this stage, the low-frequency new mutations are detected as rare minor alleles. These new alleles are also under negative selection. Given more time, the new alleles will rise in frequency through genetic drift, so that there is no longer an excess of rare minor alleles, and only a low polymorphism-to-divergence ratio remains. Third, the proportion of high-frequency derived alleles detects “recent” positive selection, within the past tens of thousands of years, enough time for beneficial derived alleles and their nearby linked variants to rise to higher frequencies than the ancestral alleles, but not enough time for them to complete selective sweeps and become fixed in the population. The underlying assumption is the infinite sites model, which is incompatible with reversion mutations. (B-C) Density plot displays the relationship between the number of divergent substitutions and the number of polymorphisms in (B) 50 kb and (C) 500 bp windows along each autosome. Windows that fall within Consensus HAQERs (green) clearly deviate from the genome-wide trend. (D-E) Unpolarized allele frequency spectra across segregating sites in 501 African individuals (1,002 alleles) within region sets on (D) hs1 or (E) hg38. HAQER sets show elevated proportions of rare minor alleles (minor allele frequency < 0.05) compared to RAND on hg38 but not on hs1. (F) Scatter plot of divergent sites per base against polymorphic sites per base for each region set, evaluated on either hg38 (red) or hs1 (blue). HAQERs show large differences in polymorphic site estimates across references, consistent with their enrichments in repetitive regions, which may be prone to variant calling misclassification. This suggests that the rare minor allele excess observed in HAQERs in hg38 may be the result of a variant calling artifact. (G) Derived allele frequency spectra across 501 African individuals (1,002 alleles) for segregating sites within region sets. The 3 HAQER sets have the highest proportions of high frequency derived alleles (derived allele frequency > 0.95). In the inset, the dotted line represents RAND. Statistical significance is assessed by pairwise chi-squared tests against RAND. (H) Mean selection parameters for each region set, inferred from the segregating sites of the 5 independent populations within the 501 African individuals. Based on our estimations of mean selection parameters, HAQERs and HARs are under positive selection, in contrast with Missense regions and UCEs, which are under negative selection. Statistical significance is assessed by pairwise t-tests against RAND (**** p < 0.0001). All analyses shown were conducted on hs1 unless otherwise stated.

**Figure S4:**
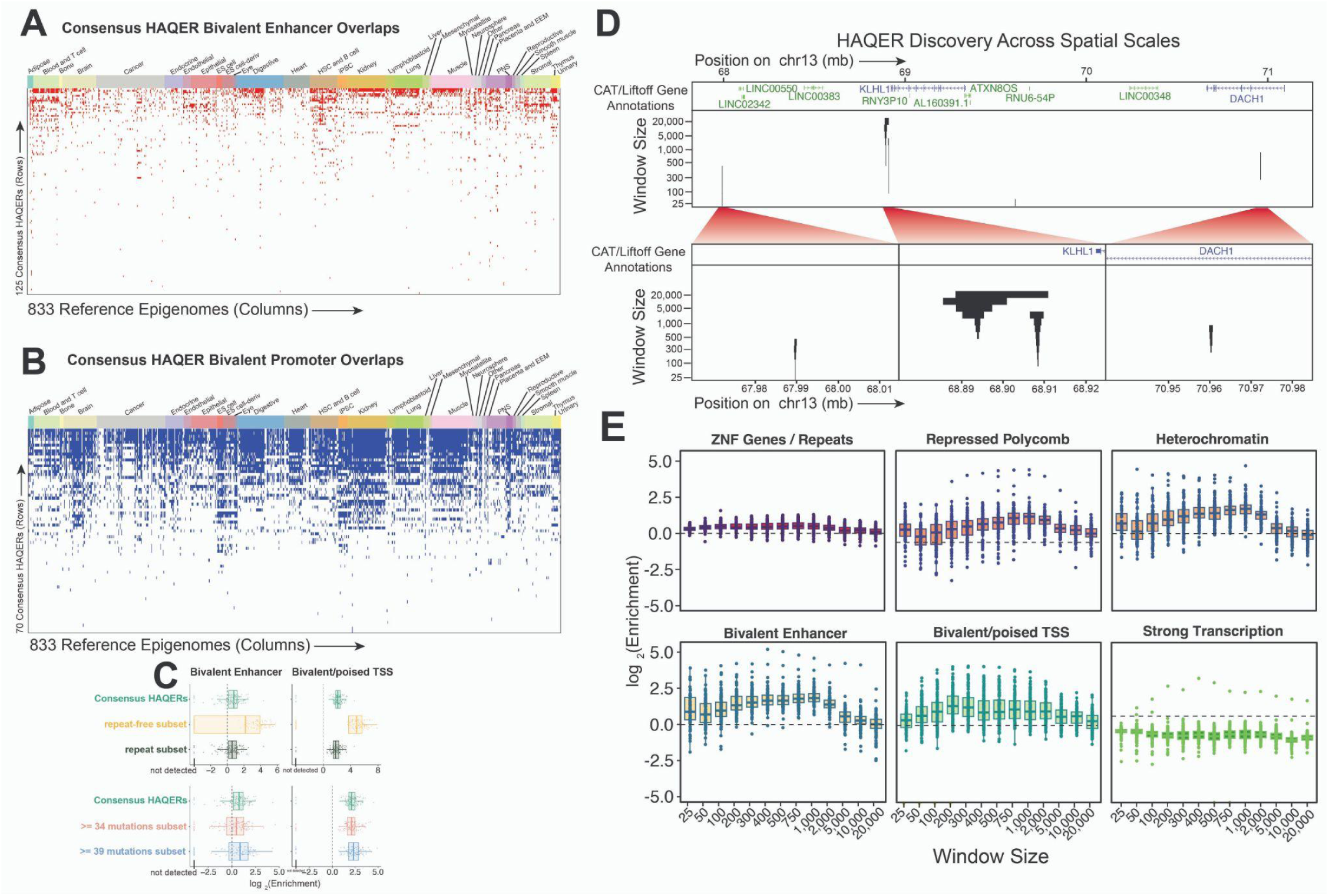
Expanded epigenomic analysis of HAQERs, and multi-scale divergence landscapes of the human genome, related to Figure 4. (A-B) Heatmap of Consensus HAQERs (rows) overlapping (B) bivalent enhancer states and (C) bivalent promoter states across 833 reference epigenomes^17^(columns). (C) Overlap enrichments between bivalent regulatory states (Bivalent Enhancer; Bivalent/poised TSS) across 127 reference epigenomes^62^ and Consensus HAQERs, including subsets restricted to HAQERs overlapping repetitive elements (repeat subset) and its complement (repeat-free subset) or subsets defined at higher divergence thresholds (≥34 or ≥39 mutations). (D) HAQER genomic locations, identified across diverse spatial scales, in a 4 mbp region of chr13. Some larger-scale HAQERs are composite elements, composed of multiple HAQER elements identified at smaller window sizes (middle). (E) Chromatin state enrichments for T2T HAQERs identified at a range of window sizes across 127 reference epigenomes^62^.

**Figure S5:**
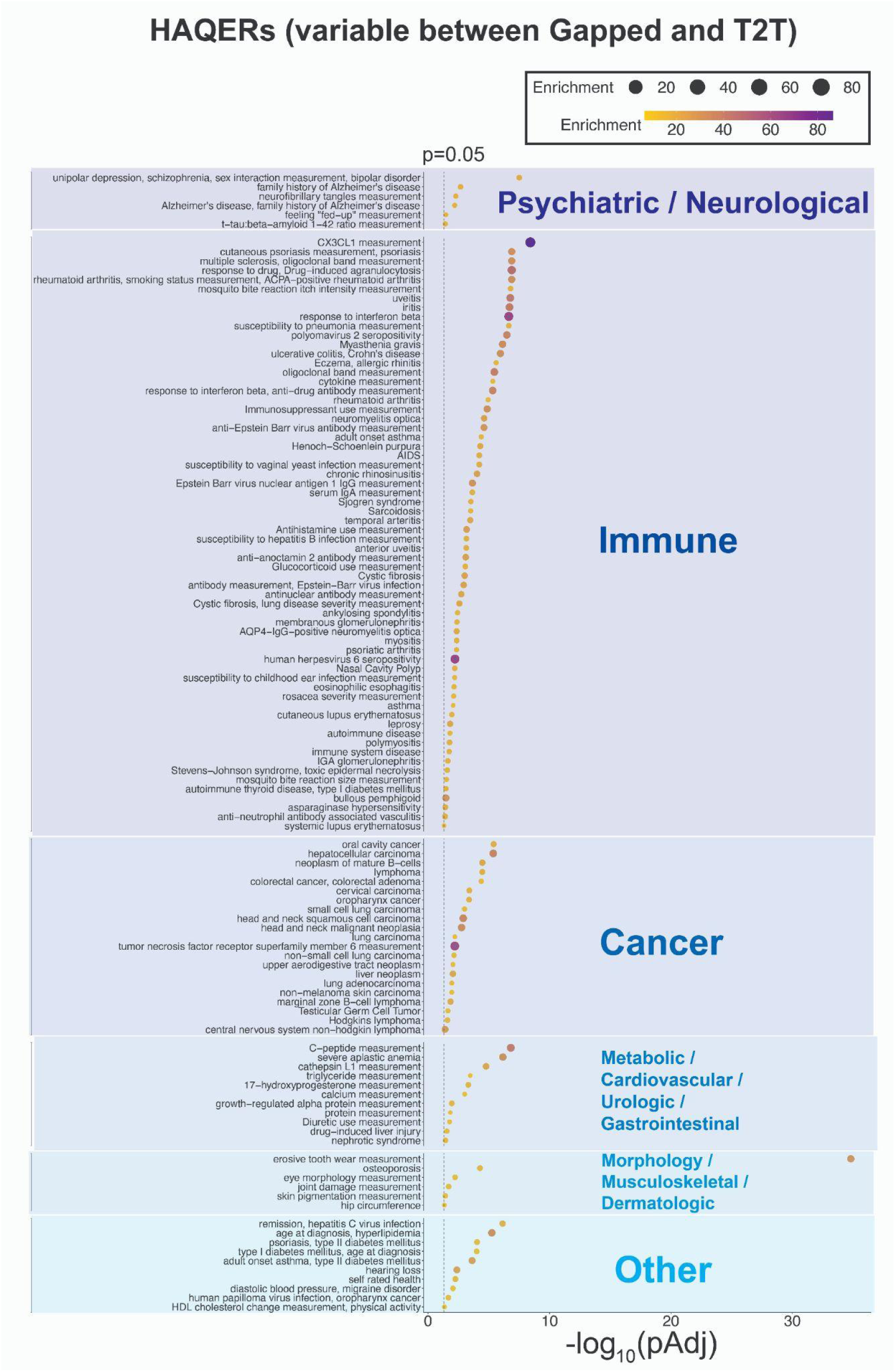
GWAS trait enrichments for variable HAQERs, related to Figure 4. Overlap enrichments between the variable HAQER set – which consists of Gapped HAQERs not overlapping T2T HAQERs and T2T HAQERs not overlapping Gapped HAQERs, and GWAS (genome-wide association study) loci, together with their linked variants (Plink R^2^ > 0.7). Traits with significant FDR-adjusted enrichments (p < 0.05) are shown. Variable HAQERs display widespread enrichments across psychiatric, neurodegenerative, immune, metabolic, and cancer-related traits.

**Figure S6:**
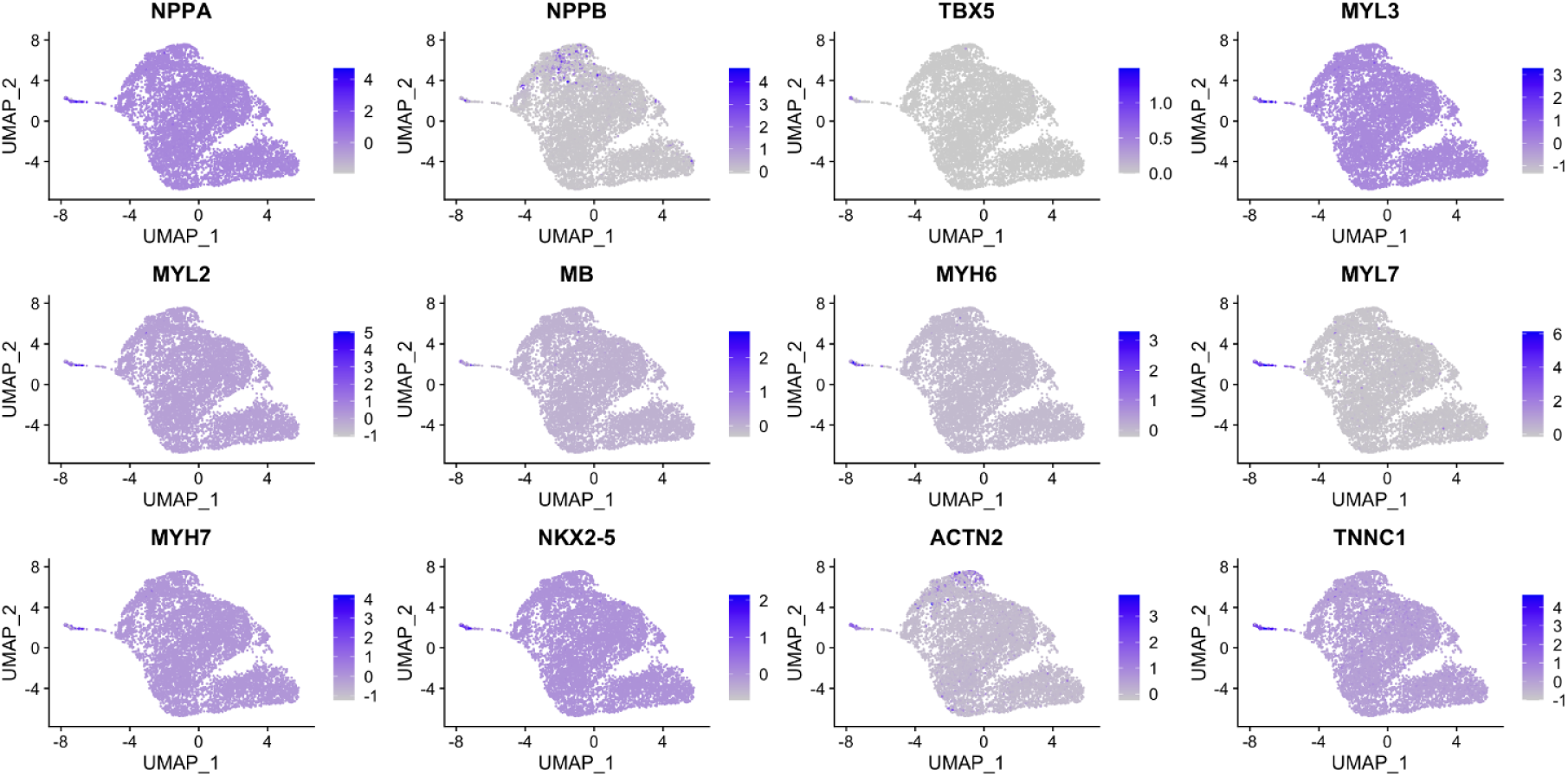
iPSC-derived cardiomyocytes express canonical cardiomyocyte marker genes, related to Figure 5. UMAP projection of iPSC-derived cardiomyocytes highlighting expression of select marker genes, including contractile machinery genes such as myosin light (*MYL2/3/6*) and heavy chains (*MYH6/7*), myoglobin (*MB*), natriuretic peptides (*NPPA* and *NPPB*), and transcriptional and developmental regulators (*NKX2-5* and *TBX5*).

